# Climate change and local host availability drive the northern range boundary in the rapid northward expansion of the eastern giant swallowtail butterfly

**DOI:** 10.1101/868125

**Authors:** J. Keaton Wilson, Nicolas Casajus, Rebecca A. Hutchinson, Kent P. McFarland, Jeremy T. Kerr, Dominique Berteaux, Maxim Larrivée, Kathleen L. Prudic

**Affiliations:** School of Natural Resources and the Environment, University of Arizona; Biology, University of Quebec Rimouski; Electrical Engineering and Computer Science, Oregon State University; Vermont Center for Ecostudies; Biology, University of Ottawa; Entomology, Montreal Space For Life

## Abstract

**Aims:** Species distributions result from both biotic and abiotic interactions across large spatial scales. The interplay of these interactions as climate changes quickly has been understudied, particularly in herbivorous insects. Here, we investigate the relative impacts these influences on the putative northern range expansion of the giant swallowtail butterfly in North America.

**Location:** North America.

**Time period:** 1959-2018.

**Major taxa studied:** Eastern Giant swallowtail, *Papilio cresphontes* (Lepidoptera: Papilionidae); common hop tree, *Ptelea trifoliata*; common prickly ash, *Zanthoxylum americanum*; southern prickly ash, *Zanthoxylum clava-herculis* (Saphidales: Rutaceae).

**Methods:** We used data from museum collections and citizen science repositories to generate species distribution models. Distribution models were built for each species over two time periods (T1 = 1959-1999; T2 = 2000-2018).

**Results:** Models for *P. cresphontes* and associated host plants had high predictive accuracy on spatially-explicit test data (AUC 0.810-0.996). Occurrence data align with model outputs, providing strong evidence for a northward range expansion in the last 19 years (T2) by *P. cresphontes*. Host plants have shifted in more complex ways, and result in a change in suitable habitat for *P. cresphontes* in its historic range. *P. cresphontes* has a northern range which now closely aligns with its most northern host plant - continued expansion northward is unlikely, and historic northern range limits were likely determined by abiotic, not biotic, factors.

**Main conclusions:** Biotic and abiotic factors have driven the rapid northern range expansion in the giant swallowtail butterfly across North America in the last 20 years. A number of bioclimatic variables are correlated with this expansion, notably an increase in mean annual temperature and minimum winter temperature. We predict a slowing of northward range expansion in the next 20-50 years as butterflies are now limited by the range of host plants, rather than abiotic factors.

## INTRODUCTION

Species distributions are known to be greatly influenced by climate (Brown *et al.*, 2016). Climate related range shifts have been and are continuing to be documented globally across taxa and systems: terrestrial (Parmesan & Yohe, 2003), marine (Poloczanska *et al.*, 2013), and aquatic (Rahel & Olden, 2008). With current changes in global climate, species range shifts (Parmesan *et al.*, 1999) and extensions in both altitude and latitude are being observed (Roth *et al.*, 2014; Kerr *et al.*, 2015). While many studies have examined the ongoing changes in climate on biodiversity and species ranges, most consider only abiotic factors in their analyses, missing the potential importance of local interspecific interactions once a species moves into a novel environment beyond its previous range (Blois *et al.*, 2013; HilleRisLambers *et al.*, 2013; Wisz *et al.*, 2013).

Several interspecific interactions are known to play important roles in shaping range boundaries including competition (Connell, 1961; Huey *et al.*, 2009; Stanton-Geddes *et al.*, 2012), mutualism (Chalcoff *et al.*, 2012; Moeller *et al.*, 2012), facilitation (Bader *et al.*, 2007; Stueve *et al.*, 2011; Ettinger & HilleRisLambers, 2017) and natural enemies (Freeman *et al.*, 2003; Speed *et al.*, 2010). When a species extends into a new local environment, there are a few main scenarios it can encounter (Holt, 2003; Urban *et al.*, 2007; Sexton *et al.*, 2009): 1) ecological conditions that are similar enough to previous conditions that there is little immediate effect on fitness and population growth rate, 2) the new local environment may possess biotic or abiotic conditions that differ from the original local environment and can accelerate (e.g. competitive or predatory release; or 3) decelerate (e.g. nutrient or nesting limitation) range expansion.

For insect herbivores, climate change can influence abundance and distribution through direct mechanisms (physiological impacts on growth, development and reproduction that impact fitness) and indirect mechanisms (impacting biotic factors such as host plant quality or predator abundance) (Bale *et al.*, 2002; Deutsch *et al.*, 2008; Kingsolver *et al.*, 2011; Robinson *et al.*, 2017). How and when climate change will affect herbivorous insect dynamics has received considerable attention generating a diversity of observed responses, especially in the pest management literature (Porter *et al.*, 1991; Cannon, 1998; Harrington *et al.*, 2001; Altieri *et al.*, 2015; Castex *et al.*, 2018). Some species are expanding in ranges and abundance (Battisti *et al.*, 2005; Robinet & Roques, 2010; Robinson *et al.*, 2017) while others are retracting and decreasing in numbers (Robinet & Roques, 2010; Zvereva *et al.*, 2016; Sánchez-Bayo & Wyckhuys, 2019). Host plant abundance and distribution play a key role in generating these patterns as herbivorous insects are often limited by larval food resources (Dempster & Pollard, 1981; Pearson & Knisley, 1985; Ylioja *et al.*, 1999). Exactly how host-availability translates into patterns of distribution, abundance, and range shifts for insect herbivores is still contentious and particularly complex when combined with direct effects on physiology (Louthan *et al.*, 2015; Lany *et al.*, 2018). Our understanding of the determinants regulating species distributions are becoming more nuanced as we begin to incorporate information on species’ dispersal capacity, population abundance trends, and climatic variables into our models (Elith & Leathwick, 2009).

In this study, we investigate the role of host availability and climatic variables on the range expansion of the giant swallowtail butterfly (Papilionidae: *Papillio cresphontes*) in northeast North America over the last 60 years (1959-2018), with an emphasis on accelerated expansion in the last 18 years. We combine evidence from raw occurrence data with a series of species distribution models for *P. cresphontes* and associated host plants to evaluate the rate and direction of range changes in relation to both abiotic and biotic factors. While other studies have incorporated biotic variables as model inputs (Bueno de Mesquita *et al.*, 2016; Palacio & Girini, 2018), our approach was to model the distribution of the insect herbivore and host plants separately and using these independent models to make *post hoc* inferences and comparisons of ranges. Because both this insect and its primary larval host plants (the common prickly ash [Rutaceae: *Zanthoxylum americanum*], southern prickly ash [Rutaceae: *Zanthoxylum clava-herculis*] and common hop tree [Rutaceae: *Ptelea trifoliata*]) are conspicuous, they are often reported in systematic biological surveys and museum collections. In this study, we bring together a combination of museum collection, survey, and citizen science data to understand how host plant availability, climate changes, and butterfly abundance are influencing the rapid expansion of an herbivorous insect. This study is one of few to demonstrate the interplay of both climate change and biotic interactions in shaping range limits while focusing on the ecologically important role of herbivore and pollinator.

## MATERIAL AND METHODS

### Study region and time interval

We focused on eastern North America (study area bounded by −94° and −65° longitude and 25° and 55° latitude) where *Papilio cresphontes* has been reported to be expanding rapidly (Finkbeiner *et al.*, 2011; Breed *et al.*, 2012) and data are readily available for both *P. cresphontes* and larval host plants, (*Zanthoxylum americanum, Zanthoxylum clava-herculis* and *Ptelea trifoliata).* Though records of *P. cresphontes* exist further west than −94°, we set this cutoff to minimize complications of misidentification and complex species boundaries with its cogener *P. rumiko.* We categorized and compared two time periods: T1 (1959-1999) representing the period prior to the beginning of the rapid range expansion and T2 (2000-2018) as the period when the rapid range expansion to the north began. This cutoff point was determined from raw occurrence data (Fig. 1).

**Figure 1.**
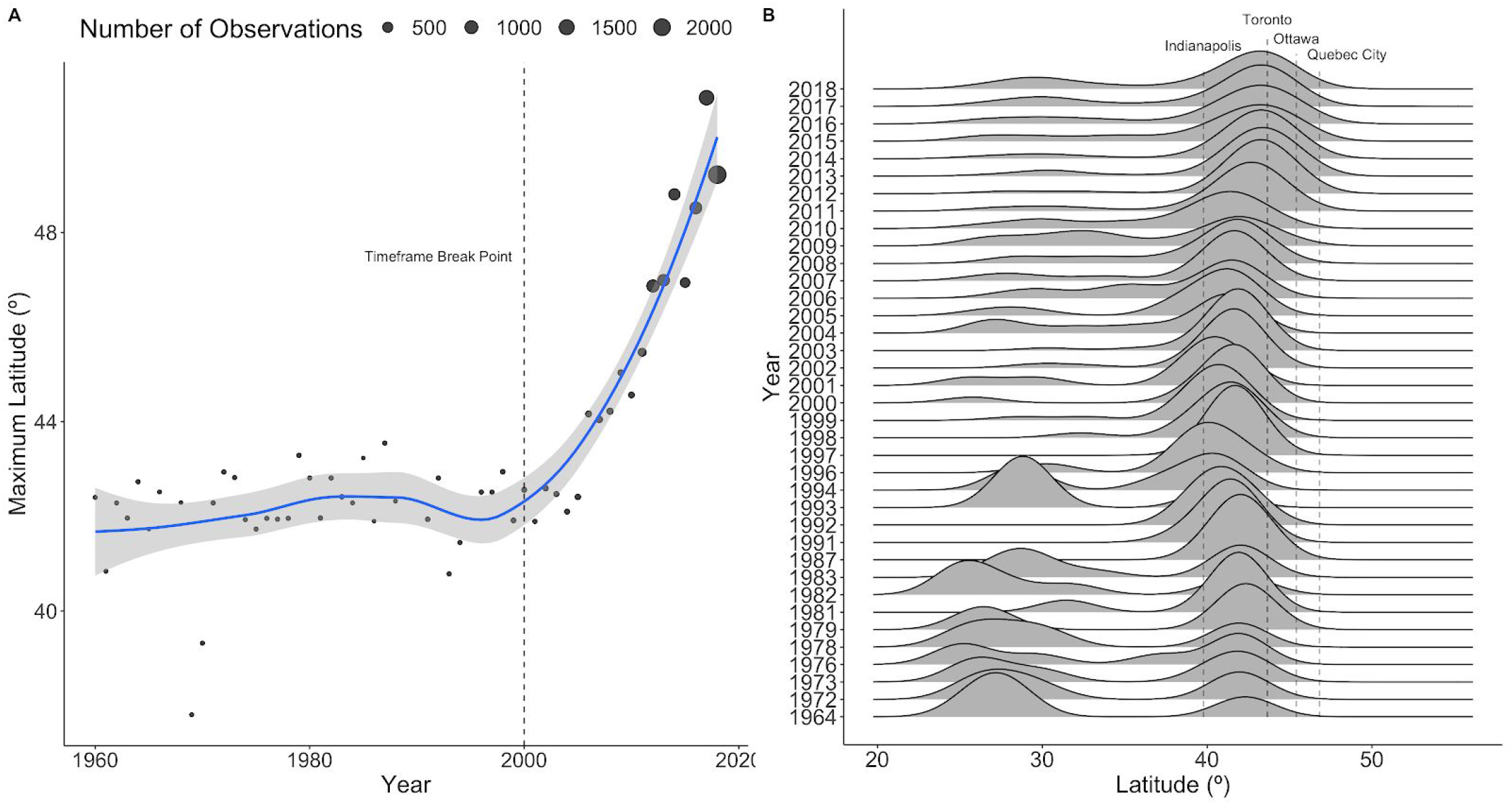
Evidence of northward range shift of *P. cresphontes* from raw occurrence data. (a) The maximum latitude of a recorded occurrence of *P. cresphontes* by year. Larger circles indicate years with higher numbers of occurrence records (high numbers in more recent years are due to increased citizen-science activity). The dashed lines represents the breakpoint between T1 and T2. Three years with extremely low maximum latitudes (< 35°) were omitted for clarity. The blue line and gray bar represent the loess smoothing curve and 95% confidence interval. (b) A ridge-plot of kernel density estimates of occurrences for *P. cresphontes.* Vertical dashed lines represent latitudes of major cities within the range. Years with < 5 occurrence records were removed from plotting.

### Data sources

#### Butterfly and host plant data

*Papilio cresphontes* (Papilionidae) is a sub-tropical butterfly widely distributed across North America. *P. cresphontes* and host plant occurrence data were obtained from a variety of sources: iNaturalist (www.inaturalist.org), n = 3,007, Global Biodiversity Information Facility (GBIF; www.gbif.org), n = 14,181, the Maine Butterfly Atlas (https://mbs.umf.maine.edu), n = 11, the Maritime Canada Butterfly Atlas (http://accdc.com/mba/index-mba.html), n = 6, Massachusetts Butterfly Club, n = 512, Butterflies and Moths of North America (www.butterfliesandmoths.org), n = 1,188, and eButterfly (www.e-butterfly.org), n = 3,083. Data from iNaturalist and GBIF were downloaded using the *spocc* package for R (Chamberlain *et al.*, 2016). We filtered iNaturalist data to include only research grade records before combining with other data sets. Combined data were filtered for time frame, duplicates, and study area extent (see below) before further analysis and model building. In total, we used 8,051 occurrence records for *P. cresphontes* and 2,697 occurrence records (combined) for all three host plant species.

#### Environmental data

We used the TerraClimate data set (Abatzoglou *et al.*, 2018), a 4 km x 4 km resolution gridded set of monthly climatological data from 1958-2017 (at the time of writing this) to generate environmental features for modeling. We calculated a set of yearly summaries of 19 bioclimatic variables (Fick & Hijmans, 2017) - frequently used in species distribution modeling) using the *dismo* package in R (Hijmans *et al.*, 2017) for each year in each time period (T1 and T2) and then averaged these summaries across each time period to provide temporally-appropriate climate summary for each set of models. We included all 19 bioclimatic variables as features during model selection and evaluation.

### Species distribution models

Distributions of *P. cresphontes* and host plants were estimated using Maxent 3.4.0, a machine learning algorithm based on the principle of maximum entropy (Phillips & Dudík, 2008; Elith *et al.*, 2011; Phillips *et al.*, 2017). Maxent is a presence-background method (Peterson et al. 2011), which is considered to perform well when modeling climatic niches across a variety of sample sizes (Wisz *et al.*, 2008). We used the *ENMevaluate* package for model building, testing, and tuning (Muscarella *et al.*, 2014), ultimately building 8 total models (*P. cresphontes* and three host species for each time period).

We used a combination of geographically-structured and regular k-fold cross validation for model testing and tuning. We generated 10,000 random background points per model and used the *blockCV* package (Valavi *et al.*, 2019a) to divide our study area into 4 km x 4 km blocks. Blocks were randomly assigned to folds 1-5 over 250 iterations to determine a block design that maximized evenness of occurence and background points spread across all folds. This procedure was repeated for every model (8 times in total). The data from folds 1-4 were used as training data for Maxent cross-validation and tuning, while fold 5 was reserved as a set of out-of-sample test data for final model evaluation. Throughout the manuscript, we refer to these data as test data. We used another set of random 5-fold cross validation within the training data while to tune model parameters. Throughout the manuscript, we refer to these data as validation data. We tested the full suite of Maxent feature combinations (linear, quadratic, product, threshold, hinge and combinations) and a combination of regularization multipliers (0.5-4 in 0.5-step increments). We examined models using a range of evaluation metrics (Supplementary Figures 1-8), but eventually chose the model with the highest area under the receiver operating characteristic curve (AUC) on validation data. All evaluation metrics were reported for the separate set of spatially-explicit test data generated by *blockCV* (Table 1). Following the classification of (Swets, 1988), AUC values range between 0.5 for models with no predictive ability and 1.0 for models giving perfect predictions; hereby, values > 0.9 describe a ‘very good’, > 0.8 a ‘good’ and > 0.7 a ‘useable’ discrimination ability of the model and allow to assess the ability of the model to distinguish between species records and background data (Phillips *et al.*, 2006). Once the optimal parameters for a given species and time-frame were determined, we built full models using all available occurrence data to generate predictions for subsequent visualizations and analyses. We mapped the ‘cloglog’ Maxent output, which can be interpreted as probability of occurrence under the assumption that the species presence or absence at nearby sites are independent ((Phillips & Dudík, 2008; Elith *et al.*, 2011; Phillips *et al.*, 2017). Importance of predictors was assessed using the percent importance metrics generated when building full models. These metrics are built as Maxent steps through modifications of coefficients for single features - Maxent tracks which environmental variables those features depends on, and calculates the total percentage of contribution of each environmental variable at the end of training (Phillips *et al.*, 2006). Thresholds for binary presence-absence maps and presence distributions were generated using the maximum test specificity plus sensitivity (Liu *et al.*, 2005).

**Table 1.** Model features and evaluation metrics on geographically-structured test data

| Species | Timeframe | Occurrences* | Feature Classes** | Regularization Multiplier | AUC (test data) | Threshold |
| --- | --- | --- | --- | --- | --- | --- |
| *P. cresphontes* | T1 | 219 | LQH | 1 | 0.924 | 0.095 |
|  | T2 | 7832 | LQH | 1 | 0.869 | 0.188 |
| *Z. americanum* | T1 | 153 | LQHP | 0.5 | 0.971 | 0.293 |
|  | T2 | 1170 | H | 2 | 0.979 | 0.266 |
| *Z. clava-herculis* | T1 | 9 | LQHP | 0.5 | 0.996 | 0.956 |
|  | T2 | 364 | LQHP | 1 | 0.892 | 0.189 |
| *P. trifoliata* | T1 | 139 | LQHP | 0.5 | 0.960 | 0.154 |
|  | T2 | 862 | LQHP | 0.5 | 0.810 | 0.187 |

\* Full number of occurrences, not the number of occurrences within the test set.
\*\*Feature classes tuned in MaxEnt (L = linear, Q = quadratic, H = hinge, P =product, T = threshold)

Maxent has become a popular modeling resource because of its predictive power, ease of use, and a well-detailed literature to get researchers started (Phillips & Dudík, 2008; Elith *et al.*, 2011; Phillips *et al.*, 2017). However, this framework has also received criticism, with researchers advocating for more explicit examinations of tuning parameters, evaluation metrics, and the incorporation of tools to deal with sampling bias (Radosavljevic & Anderson, 2014). Recent software additions have addressed some of these challenges, and opened up the ‘black-box’ of Maxent (Phillips *et al.*, 2017), though issues remain, particularly in the transparency of researchers’ hyperparameter tuning and evaluation (Morales *et al.*, 2017). To this end, we have implemented recently developed tools *(ENMevaluate* and *blockCV* packages in R; (Muscarella *et al.*, 2014; Valavi *et al.*, 2019b)) to explicitly outline tuning (Supplementary Figures 1-8), and to incorporate a spatially-independent evaluation design to minimize overfitting (along with the built-in regularization in Maxent).

### Northern Range Limits

We calculated the distance between the northern limit modeled for *P. cresphontes* for T1 and T2 using a longitude class approach (Leroux *et al.*, 2013). For each 4-km longitude class (i.e. the horizontal resolution of the grid used to predict species range), we determined the latitude of the northernmost grid cell where the species was predicted to be present during T1 and T2. We selected the latitude-pairs (pairs of data for a single latitude at T1 and T2) for which we had grid cells with occurrence for *P. cresphontes* in both time periods for each longitude class and tested whether the average northern limit distribution of *P. cresphontes* differed between T1 and T2, using a non-parametric paired t-test. We used similar methods to determine differences between northern range limits of *P. cresphontes* and *Z. americanum* for both time periods. All analyses were performed using the R statistical software version 3.6.0 (R Core Team 2019).

## RESULTS

### Evidence of northward range shift of *P. cresphontes* from raw occurrence data

Patterns of occurrence (as opposed to the predictive outputs from species distribution models) indicate a strong trend of a rapid and recent northward range expansion in *P. cresphontes* since the earliest recorded records of the species in our dataset (1959). The butterfly’s highest recorded latitude in a given year has increased dramatically since 2000 (Figure 1a), and the likelihood of occurrence has shifted from rare to frequent in many cities close to the current northern edge of the range (Figure 1b).

### Predictive accuracy of species distribution models

Hyperparameter tuning was performed using a variety of evaluation metrics (Supplementary Fig. 1-8), but ultimately, the feature classes and regularization multiplier of the final model used for each species-time period pair were chosen because they produced the highest AUC score on validation data (within Maxent, opposed to the spatially-explicit test data we used for final evaluation). Once the final feature set was chosen, models were evaluated on spatially-explicit out-of-sample test data created by *blockCV*. Overall, models had high predictive accuracy on test data, with AUC scores ranging from 0.810 to 0.996 (Table 1). Final models were generated using the parameter set (feature classes and regularization multiplier) described above, but built with the full set of data (training + test) to generate predictive maps (Fig. 2-3) and distributions (Fig. 4-5).

**Figure 2.**
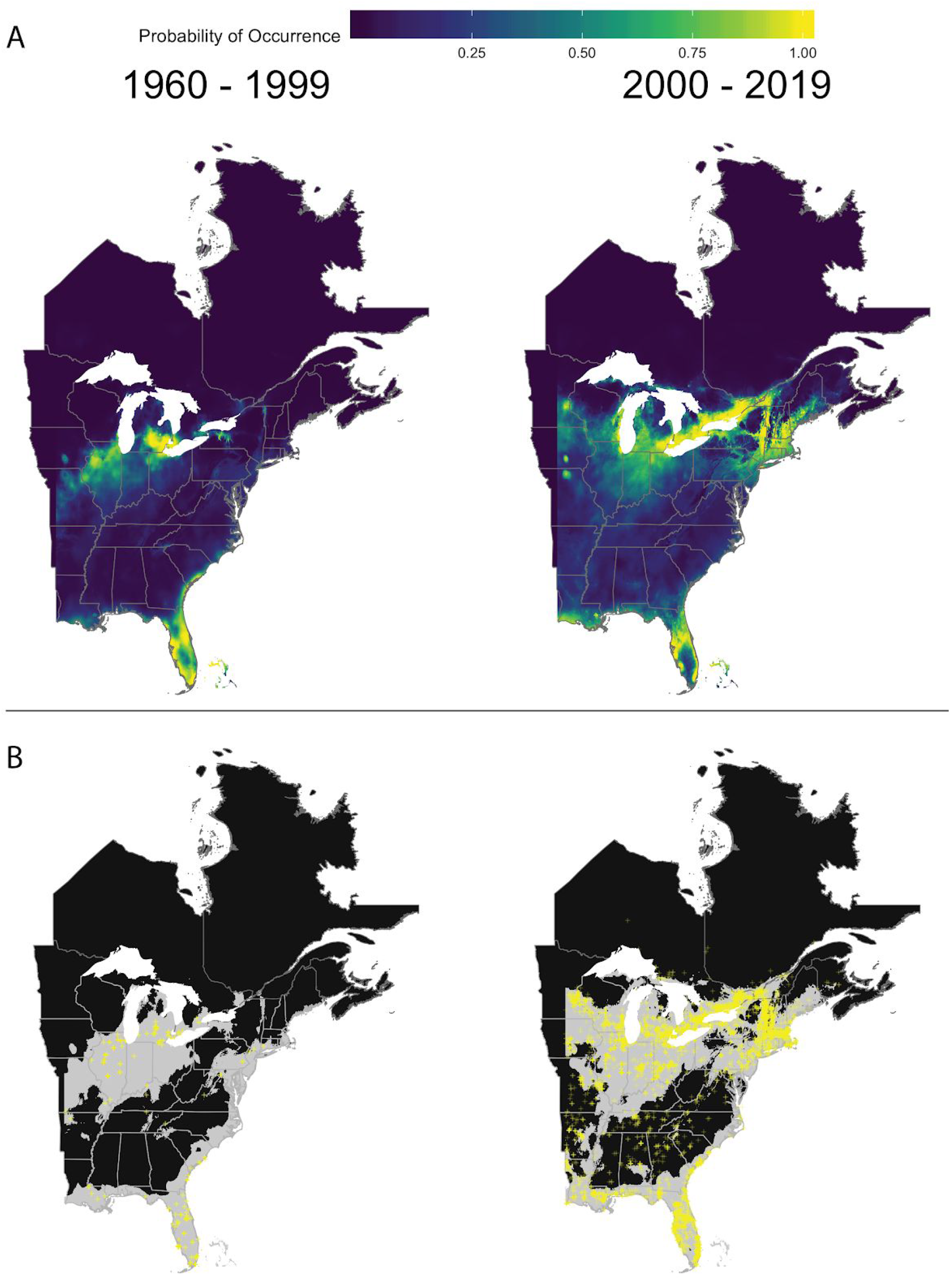
Maxent model predictions for *P. cresphontes* for t_1_, (1959-1999) and t_2_ (2000-2018). (a) cloglog transformed output from full MaxEnt models for two time periods. Lighter yellow areas denote higher probabilities of occurrence. (b) Threshold maps of presence absence for the two time periods. Gray areas represent predicted occurrence and black represents predicted absence. Yellow crosses represent actual occurrence data for each time period.

**Figure 3.**
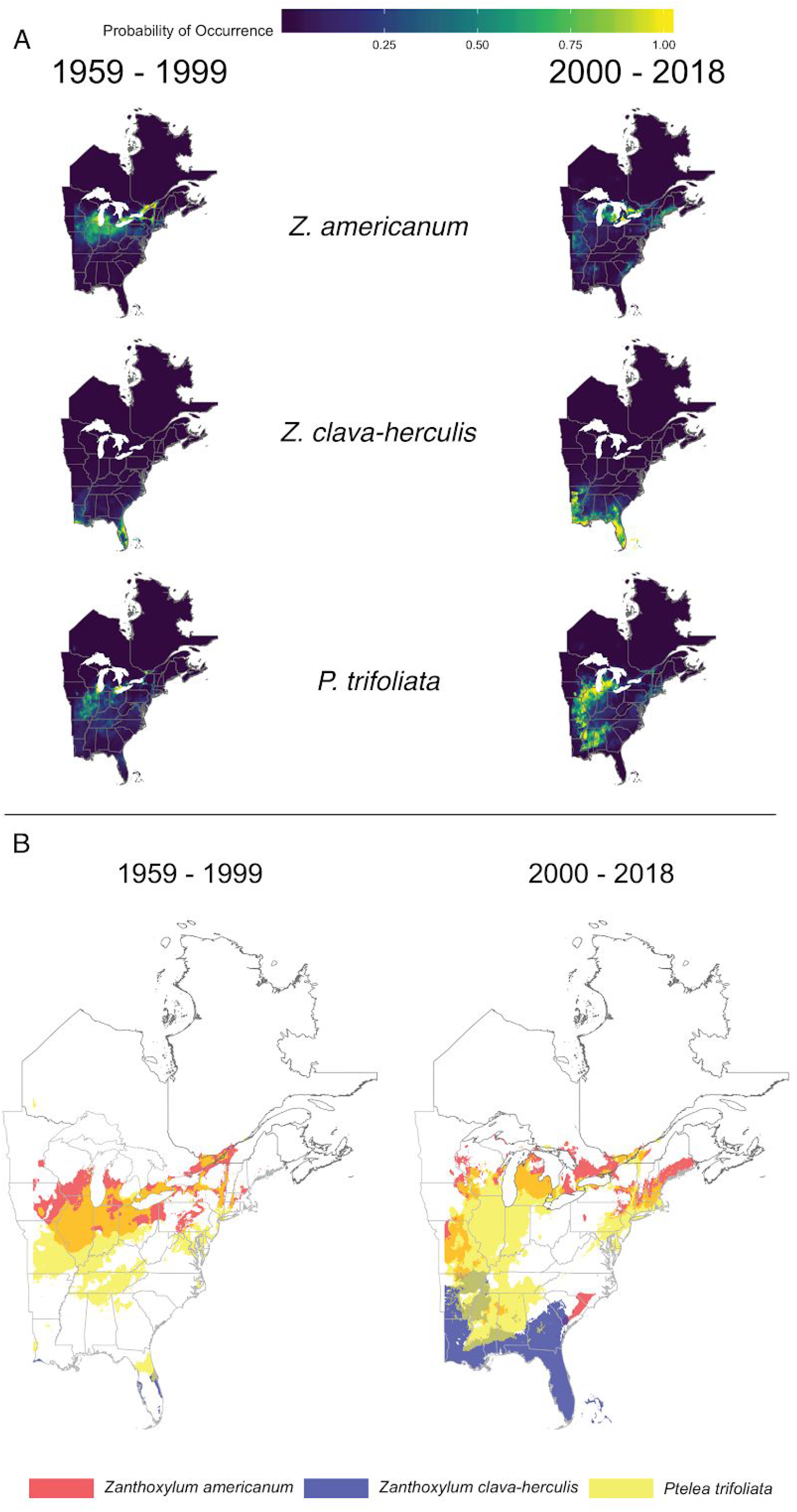
Maxent model predictions for host plants (*Z. americanum, Z. clava-herculis* and *P. trifoliata*) for t_1_ (1959-1999) and t_2_ (2000-2018). (a) cloglog transformed output from full maxent models for each host plant across two time periods. Lighter yellow areas denote higher probabilities of occurrence. (b) Threshold maps of presence absence for the two time periods. Different colors (red, blue, yellow) represent areas of predicted occurrence for each host plant and black represents predicted absence. Mixed colors (orange, purple, brown) indicate areas of overlap.

**Figure 4.**
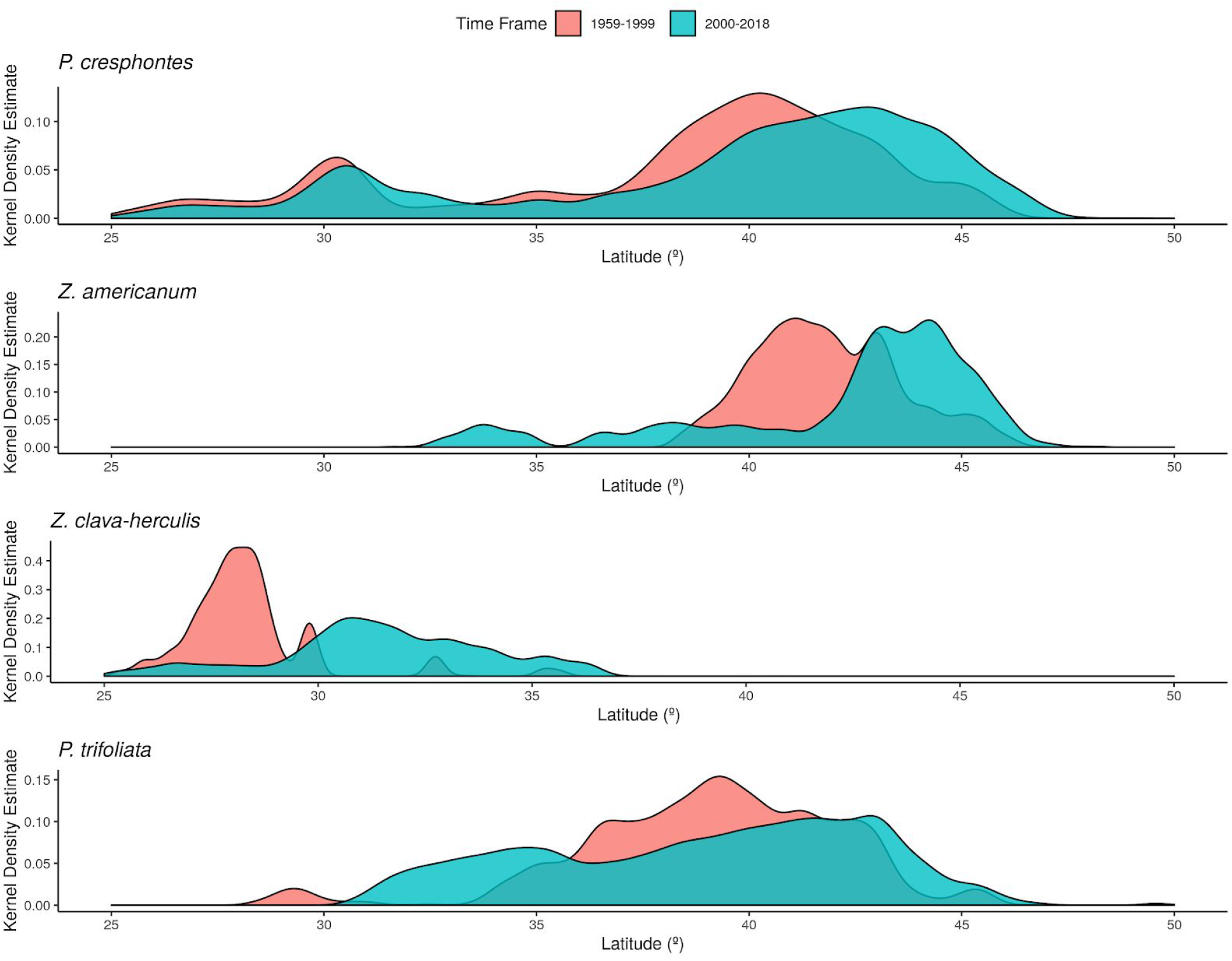
Kernel density estimates for modeled predicted presence of *P. cresphontes* and host plants (*Z. americanum, Z. clava-herculis* and *P. trifoliata*) for T1 (1959-1999) and T2 (2000-2018). Red plots are from T1 (1959-1999) and blue plots are from T2 (2000-2018).

**Figure 5.**
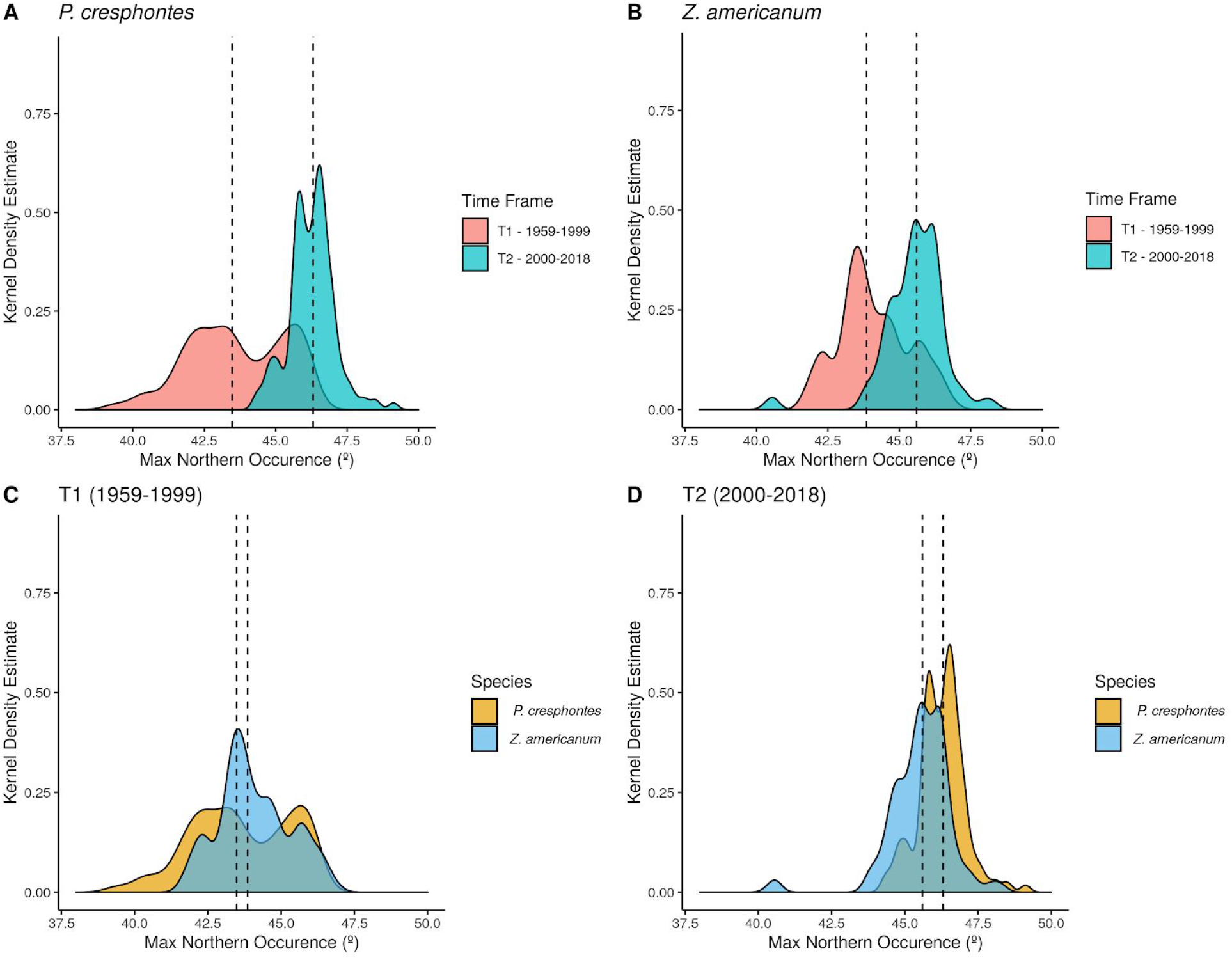
Predicted maximum northern occurence for *P. cresphontes* and host plants for T1 (1959-1999) and T2 (2000-2018). Dashed vertical lines represent the median value for each group, (a) *P. cresphontes* northern range limit between the two time periods. (b) *Z. americanum* northern range limit between the two time periods. (c) Northern range limit comparison for T1 (1959-1999) for *P. cresphontes* and *Z. americanum*. (d) Northern range limit comparison for T2 (2000-2018) for *P. cresphontes* and *Z. americanum*.

### *Papilio cresphontes* has expanded northward due to recent climate warming

Predictive maps generated from MaxEnt models clearly show a change in the distribution of *P. cresphontes* between T1 and T2, with a northward expansion since 2000 (Fig. 2a-b). Kernel density estimate plots generated from threshold occurence predictions mirror this result (Fig. 4), and highlight that different parts of *P. cresphontes’* range match host plant use. *Z. americanum* closely matches *P. cresphontes* in the north, while the middle and southern part of the range is defined by the presence of *Z. clava-herculis* and *P. trifoliata*.

### Host plant range shifts

Overall, host plants (*Z. americanum, Z. clava-herculis* and *P. trifoliata*) demonstrated more complex changes in distribution between T1 and T2 compared to *P. cresphontes* (Fig. 3a-b). Historically, the species were split latitudinally (with significant overlap) with *Z. americanum* occupying the northern part of the study area, *P. trifoliata* the middle, and *Z. clava-herculis* in the far south (Fig 3b). However, this pattern changes in T2, with a range expansion of *Z. americanum* northward, but also westward to the boundary of our study area. There are also significant distribution changes for the other two host species, with *Z. clava-herculis* expanding northward out of Florida and across the southeast coastal areas and *P. trifoliata* expanding southward throughout the southeastern United States.

### Northern range limits for *P. cresphontes* have shifted northward and closely match *Z. americanum*

The northern range limit of *P. cresphontes* was significantly higher in T2 compared to T1 (t = −35.08, df = 563, p < 0.001; Fig. 5a) where the average northern-most occurence for T2 (median = 46.3125 ± 0.937°) was 2.83° (~ 311 km) higher in latitude than T1 (median = 43.47917 ± 1.699°). *Z. americanum* also demonstrated a significant northern range shift between T1 and T2 (t = −19.302, df = 559, p < 0.001; Fig. 5b) where the average northern-most occurence for T2 (median = 45.604 ± 1.261°) was 1.75° (~ 192 km) higher in latitude than T1 (median = 43.85417. ± 1.217°). We also tested whether the northern range limits of *P. cresphontes* and *Z. americanum* differed from each other during each time period. In each time period, there was a significant difference between the northern range limits of *P. cresphontes* and *Z. americanum* (T1: t = −8.3712, df = 506, p < 0.001; T2: t = 13.014, df = 635, p < 0.001). The difference between butterfly and host plant northern range limits shrank from 0.75° (~ 82 km) in T1 (with *Z. americanum* having a higher northern range limit) to 0.71° (~ 78 km) in T2 (with *P. cresphontes* having a slightly higher northern range limit; Fig 5b-c).

### Climatic variation in the study area between T1 and T2

Overall, T2 had a higher mean annual temperature (9.45 ± 6.20° C) than T1 (8.67 ± 6.27° C)(t = −45.274, df = 534850, p < 0.001). Bioclim variables 6 (minimum temperature of the coldest month), 8 (mean temperature of the wettest quarter), and 10 (mean temperature of the warmest quarter) had the largest importance in predicting the occurrence of *P. cresphontes* and host plants (Figure 6). These variables showed significant differences between T1 and T2 on average across our study area, with an overall trend of warmer patterns from 2000-2015 (T2) compared to 1959-1999 (T1) (Table 2).

**Figure 6.**
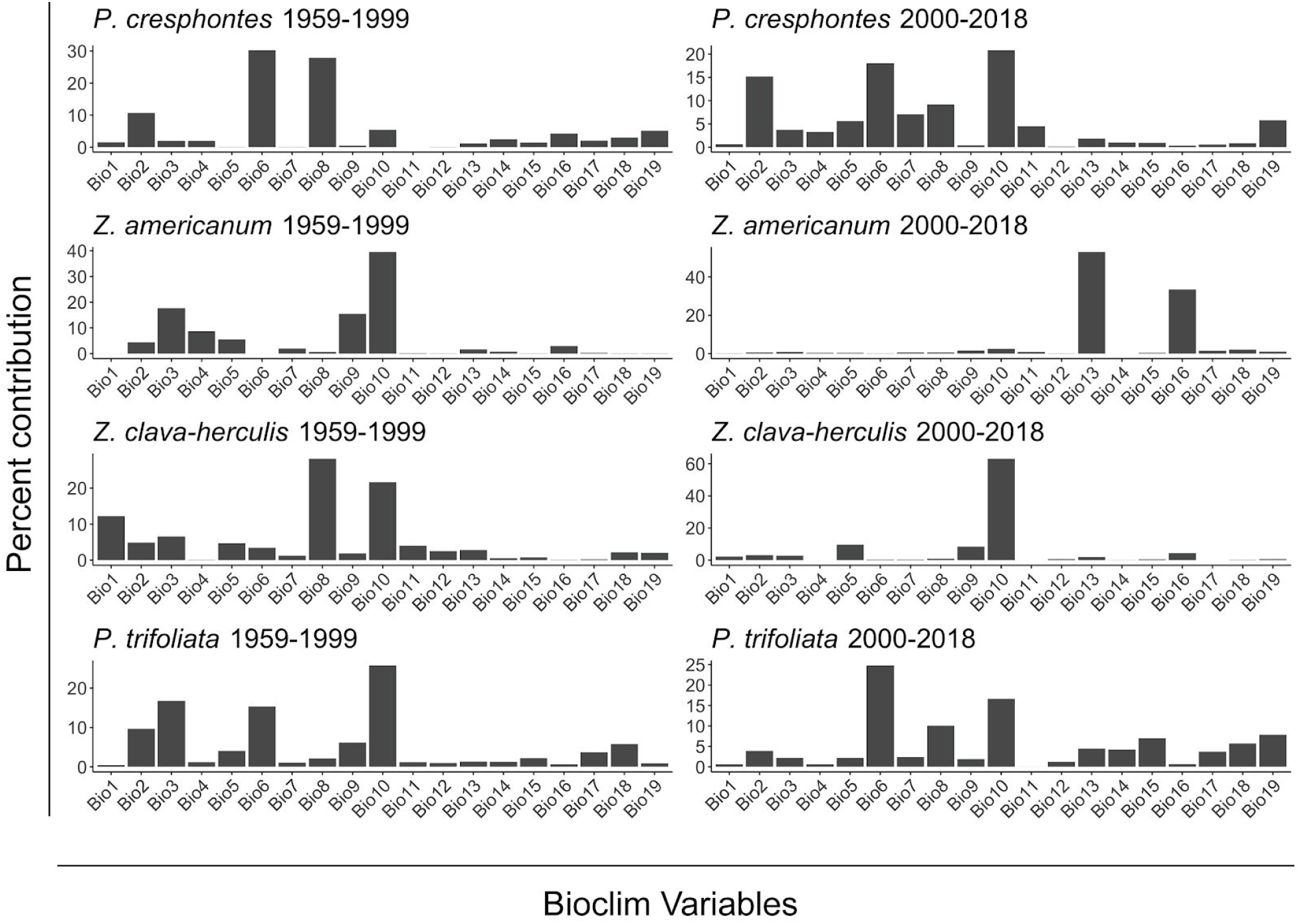
Percent contribution of each of the 19 bioclim variables to final models for each species and time period. Percentages are computed from Maxent model training - as predictive gains increase, environmental factors contributing to feature generation are calculated and summarized in the final model. Common major contributors across many models include BIO6(minimum temperature of the coldest month), BIO8 (mean temperature of the wettest quarter) and BIO10 (mean temperature of the warmest quarter).

**Table 2.**
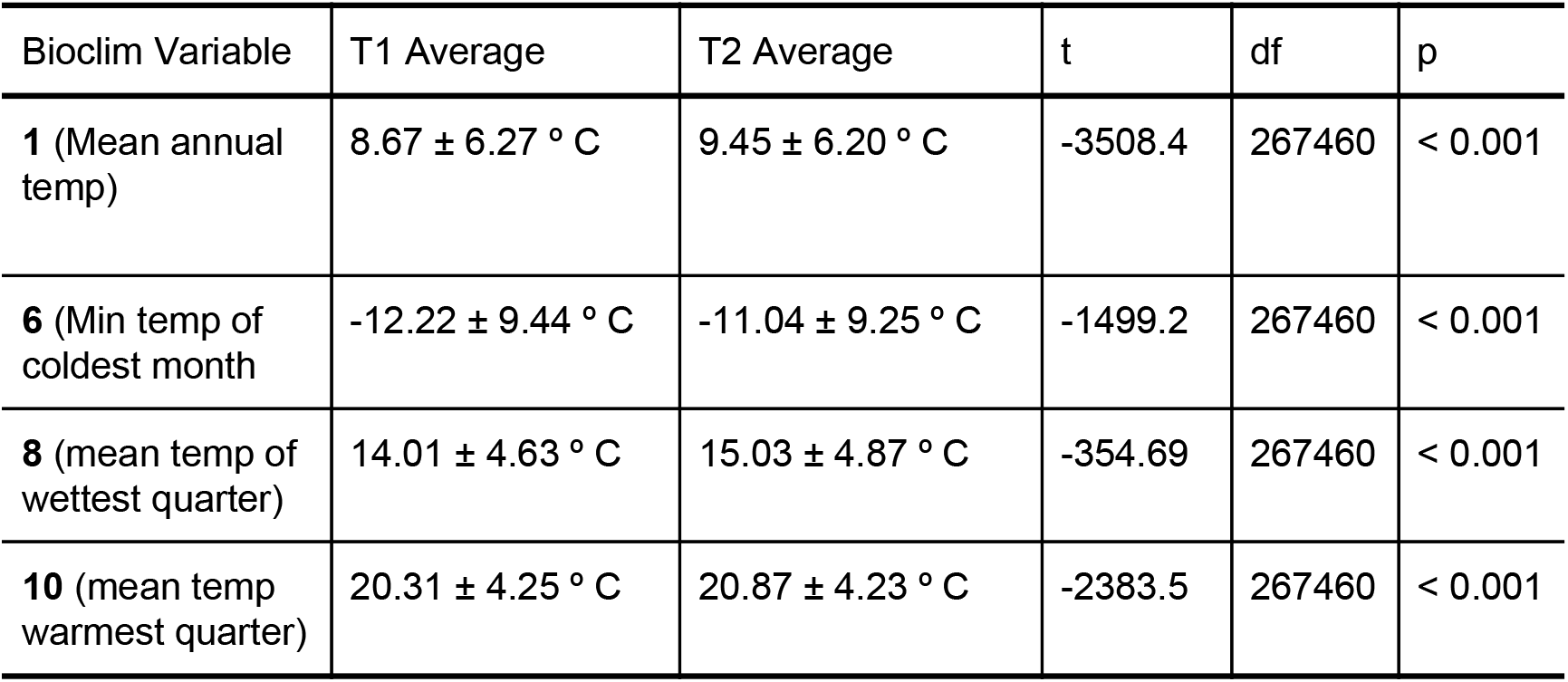
Bioclimatic shifts between T1 (1959-1999) and T2 (2000-2015)

| Bioclim Variable | T1 Average | T2 Average | t | df | p |
| --- | --- | --- | --- | --- | --- |
| **1** (Mean annual temp) | 8.67 ± 6.27 ° C | 9.45 ± 6.20 ° C | -3508.4 | 267460 | < 0.001 |
| **6** (Min temp of coldest month) | -12.22 ± 9.44 ° C | -11.04 ± 9.25 ° C | -1499.2 | 267460 | < 0.001 |
| **8** (mean temp of wettest quarter) | 14.01 ± 4.63 ° C | 15.03 ± 4.87 ° C | -354.69 | 267460 | < 0.001 |
| **10** (mean temp warmest quarter) | 20.31 ± 4.25 ° C | 20.87 ± 4.23 ° C | -2383.5 | 267460 | < 0.001 |

## DISCUSSION

The determinants of species distributions have long been debated not just because they are essential in ecology and evolutionary biology, but also because where organisms are and where they will be is central to successful conservation and restoration practices in light of rapid climate change (Buckley *et al.*, 2013; Gallagher *et al.*, 2013; Robillard *et al.*, 2015). Our study details a recent and rapid northward range expansion by *P. cresphontes* between 2000 and 2018 (Fig. 1). We also model the distributions of the butterfly’s naturally occurring larval host plants, which, when combined with analysis of *P. cresphontes* range, result in different predictions for the future distribution of this butterfly than if we had relied on abiotic variables alone (Fig. 2-3). Recent climatic shifts have allowed *P. cresphontes* to rapidly expand northward to now match or even surpass the slower moving northward range expansion of the northernmost host plant, *Z. americanum*, with further northward expansion of *P. cresphontes* now limited by host plant range (Fig. 4). Our results highlight the importance of including biotic interactions (and interactions between herbivorous insects and host plants in particular) in examinations of range shifts, an idea often highlighted, (Urban *et al.*, 2016) but infrequently implemented (Lemoine, 2015; Dilts *et al.*, 2019; Svancara *et al.*, 2019).

Poleward range shifts in herbivorous insects, particularly butterflies, have been documented for a number of species (Parmesan *et al.*, 1999; Warren *et al.*, 2001; Pöyry *et al.*, 2009; Breed *et al.*, 2012). Additionally, northward expansions of other butterfly species have been shown to have dramatic impacts on community composition through linked biotic interactions (Audusseau *et al.*, 2017), which could be happening in this system as well, but would require further examination to determine. While studies demonstrating range shifts in multiple taxa provide valuable insights into the magnitude and direction of shifts for different taxa, gaps in knowledge remain (Pöyry *et al.*, 2009). Namely, 1) how has warming acceleration affected recent range shifts during the last 10-15 years in poleward latitudes, and 2) how do abiotic and biotic factors interact to shape range shifts? Our study addresses both of these questions. We show a rapid northward range shift in *P. cresphontes* over the last 18 years (predicted most northward occurrences differ by 2.83° of latitude (~ 311 km) between T1 and T2, or a northward expansion of 165 km/decade) that is more than 27 times faster than the average of northward movement of global meta-analyses for plants, lichens, birds, mammals, insects, reptiles and amphibians, fish and marine organisms (Parmesan & Yohe, 2003) and over 9 times faster than butterfly species in Britain (Hickling *et al.*, 2006). Our findings largely follow Pöyry *et al.* (2009) who postulate that mobile species utilizing woody host plants should exhibit large range shifts northward, and that habitat availability and dispersal capacity largely determine success.

Interestingly, the northward incursion of *P. cresphontes* in northeastern North America is not a new phenomenon. Accounts detail movement into the region 145 years ago that lasted several decades (Scudder 1889). In 1875, *P. cresphontes were* found in southern New England and by 1882 there are documented records just south of Montreal, Quebec. By the 1930s, the species had apparently retracted southward and were considered ‘extremely rare’ in Massachusetts (Farquhar 1934) and didn’t push northward into the region again until recently. Multiple long-term climate reconstructions (paired with historic instrument data) for the 145-year incursion period indicate a strong warming trend compared to the previous century (Marlon *et al.*, 2016). However, this warming trend continues through the 1930s, so it is unclear which factors may have resulted in a retraction, though hydroclimatic reconstructions indicate an increase in drought in the northeastern United States over this time period, which likely had strong impacts on vegetation and host-plant distributions and quality through the range of *P. cresphontes (Marlon *et al.*, 2016).*

Our work also highlights the importance of including biotic interactions when predicting range shifts. *P. cresphontes’* current northern range now closely matches the northernmost host plant (*Z. americanum*) (Fig. 3, 5d) and is now limited by the ability of *Z. americanum* to expand its range northward. Because of the differences in life-history strategies, dispersal capabilities, reproductive outputs and environmental tolerances between *P. cresphontes* and *Z. americanum*, the northern expansion of *P. cresphontes* appears to now be largely curbed. Though sightings of the mobile adult stage of *P. cresphontes* will likely continue to be seen further north than the naturally occurring host plant range (Fig. 5d), without a suitable host plant, further northward expansion seems unlikely in nature outside of recently documented *P. cresphontes* occurrences in horticultural settings. Horticultural settings may allow it to expand beyond the northern limits or range gaps of native host plants. *Papilio cresphontes* lay eggs and larvae feed successfully on two non-native garden plants, Garden Rue (*Ruta graveolens)* and Gas Plant or Dittany (*Dictamnus albus*). It also uses Wafer Ash or Common Hop Tree *(Ptelea trifoliata)*, which is planted as an ornamental in the Northeast but is a native species from central and southeastern North America. We even documented the successful use of a potted, cultivated non-native *Citrus* tree kept outdoors during the growing season north of the butterfly’s range. Although these exotic plant species are not distributed uniformly across the region, dispersing *P. cresphontes* has an uncanny ability to find these host plants, perhaps further enabling it to expand its range as abiotic conditions allow. Data from citizen science sources continue to grow as platforms become more popular, and can provide tremendous boons to researchers across disciplines (Bonney *et al.*, 2009, 2014; Dickinson *et al.*, 2010), including those interested in creating species distribution models (Kéry *et al.*, 2010; Yu *et al.*, 2010). There has been debate about the quality and veracity of citizen science data, but recent work has demonstrated that citizen science initiatives can reliably produce research quality data though it often has similar biases to professionally-gathered data (Kosmala *et al.*, 2016). Here, we use citizen science data sources supplemented by data from museum collections to generate species distribution models using the well-established Maxent modeling framework (Phillips & Dudík, 2008; Elith *et al.*, 2011; Phillips *et al.*, 2017), and advocate for continued development and use of citizen science data and its pairing with museum collection data in developing species distribution models in ecology and conservation.

Though we focused mostly on the distributional changes of *P. cresphontes*, there were also surprisingly large range shifts in host plant species (Fig 3). In contrast to the straightforward northward expansion of *P. cresphontes*, the distributional changes in host plants was more complex and nuanced. *Z. americanum* and *P. trifoliata* have both shifted northward since 2000 in slightly different patterns (Fig. 3-4). While *Z. americanum* appears to have shifted mostly northward, *Ptelea trifoliata* has undergone a northward and westward shift, and occupies areas that overlap with the range of *Z. americanum* (Figure 3). The potential effects of this overlap on *P. cresphontes* (i.e., population dynamics, apparent competition, selection for oviposition behavior) are to our knowledge currently unknown, but would be interesting to examine in light of *P. cresphontes* westward expansion and previous work demonstrating significant within-population variation in oviposition behavior in *Papilio* (Thompson 1988). Interestingly, precipitation in the wettest month and in the wettest quarter (Bioclim variables 13 and 16) had the strongest impact in predicting the distribution of *Z. americanum* in T2, highlighting the impact that rainfall may have in shaping and limiting current distribution. In contrast, the range of *Z. clava-herculis* appears to have expanded in the southern United States - a large expansion from a relatively small range along the coast of the southeastern United States. Compared to pre-2000 distributions, available host plants to *P. cresphontes* are more widely distributed with greater overlap, but with notable gaps throughout Kentucky, Ohio, and Indiana. These complex distributional changes are likely driving part of the overall range shift northward for *P. cresphontes* (Fig. 1b) and could also be potential drivers of speciation, and the evolution of specialization or host plant switching (Descombes *et al.*, 2016).

## CONCLUSION

Multiple biotic interactions have evolved between insects and other species to create a wide variety of ecosystem services important to human health and wealth (Losey & Vaughan, 2006). Anthropogenic climate change and habitat loss are creating a growing urgency for quantifying range size, understanding range boundaries, and assessing range shifts across insect species in order to preserve the integrity of future ecosystem function. Our work outlines the power of using increasingly abundant citizen science data, as well as the importance of including biotic interactions alongside environmental factors when developing predictions for range shifts due to climate change. We advocate for expanded incorporation of these factors, particularly when examining insect herbivores.

## ACKNOWLEDGEMENTS

We would like to thank our reviewers and editors and Jeff Oliver for helpful comments and suggestions on the manuscript and analyses. Specifically, we’d like to thank the Global Biodiversity Information Facility, the Maine Butterfly Atlas, the Maritime Canada Butterfly Atlas, the Massachusetts Butterfly Club, Butterflies and Moths of North America and eButterfly for the use of their data. We’d also like to thank Scott Laurie and Ken-ichi Ueda at iNaturalist for their web platform, and all those relentless volunteers in various citizen science programs, past and present, who contributed these data for the greater good.

## Supplemental Figures

**Sup. Fig. 1:**
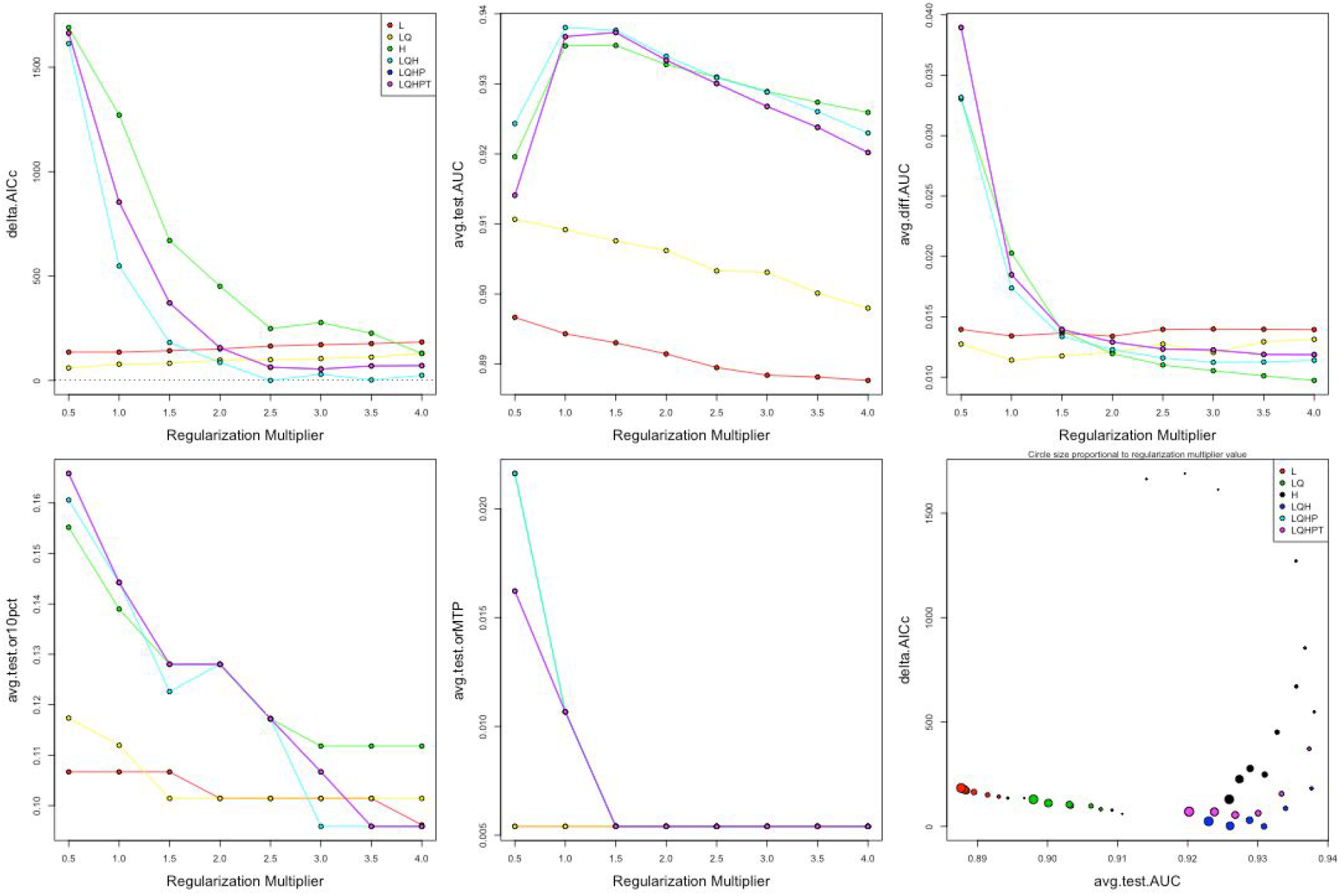
Hyperparameter tuning and model evaluation of *P. cresphontes* distribution during the T1 period.

**Sup. Fig. 2:**
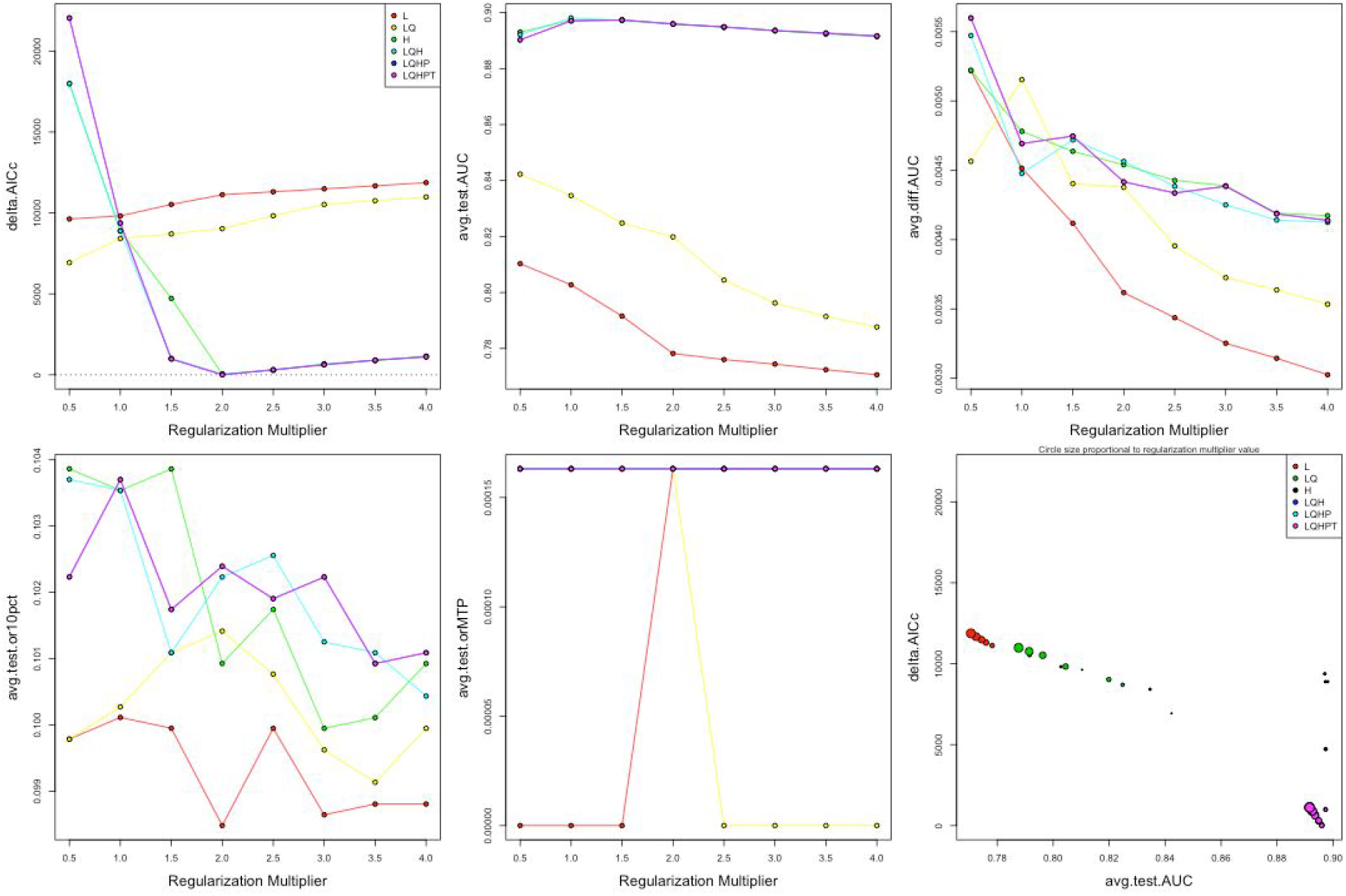
Hyperparameter tuning and model evaluation of *P. cresphontes* distribution during the T2 period.

**Sup. Fig. 3:**
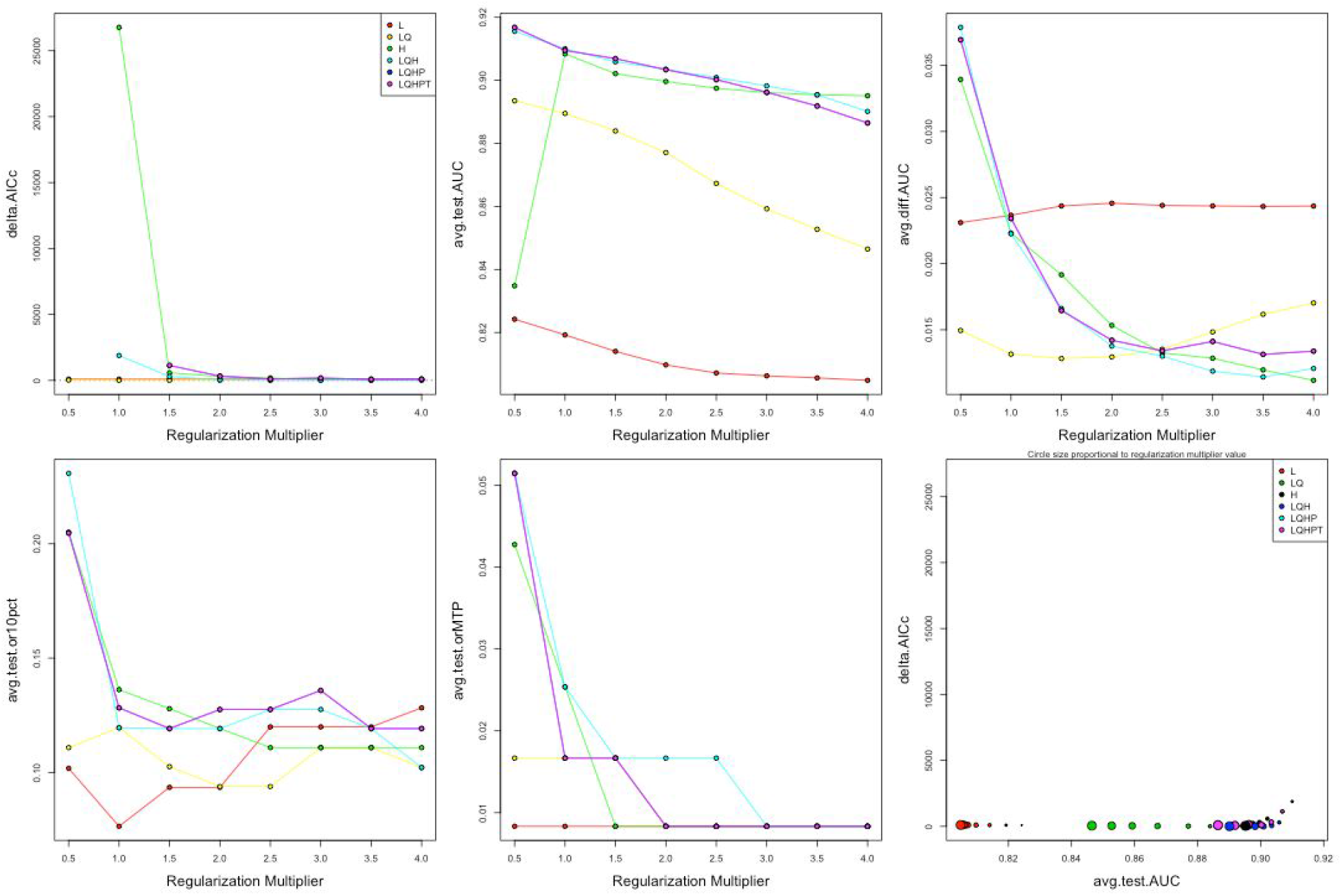
Hyperparameter tuning and model evaluation of *Z. americanum* distribution during the T1 period.

**Sup. Fig. 4:**
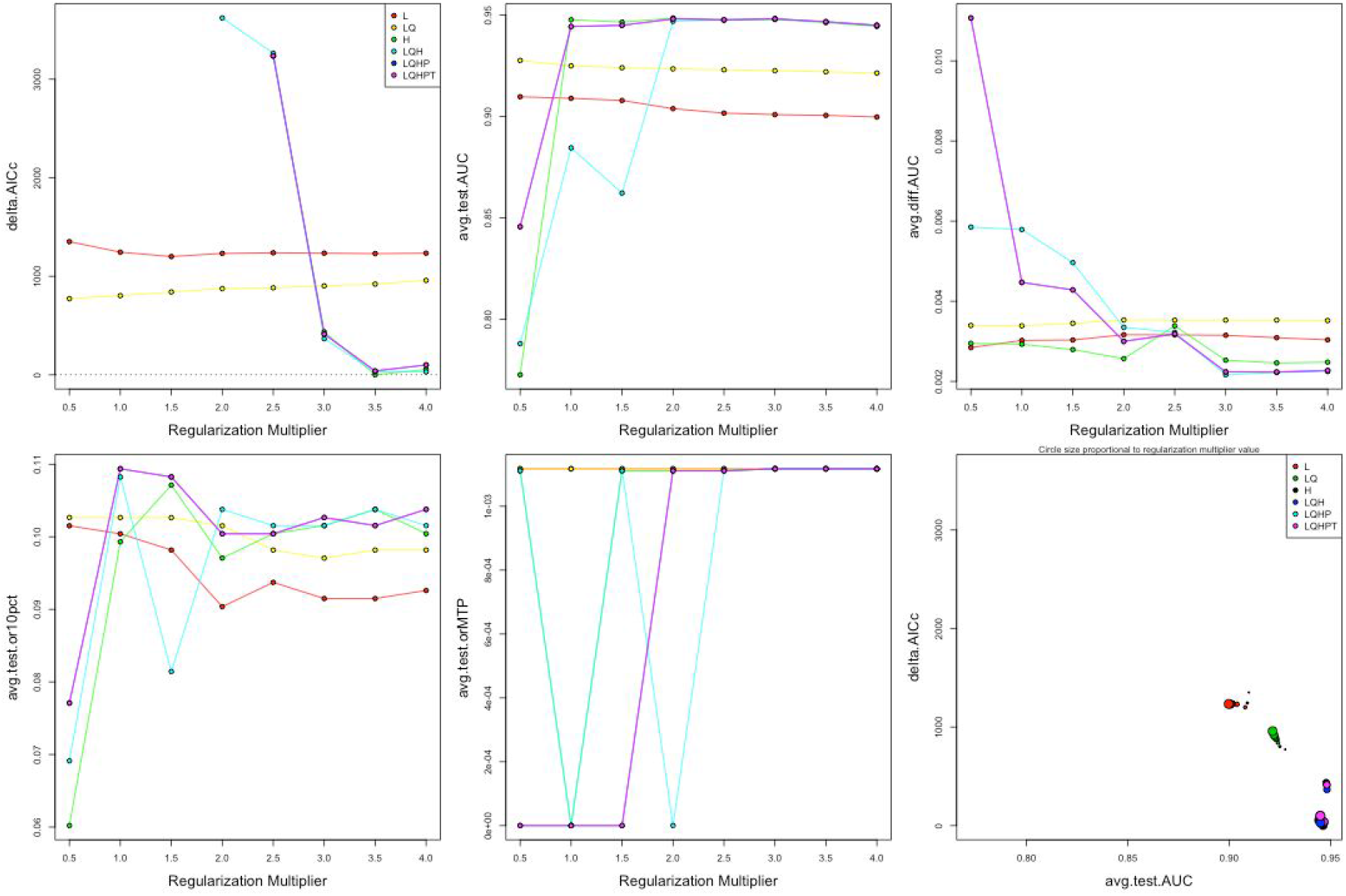
Hyperparameter tuning and model evaluation of *Z. americanum* distribution during the T2 period.

**Sup. Fig. 5:**
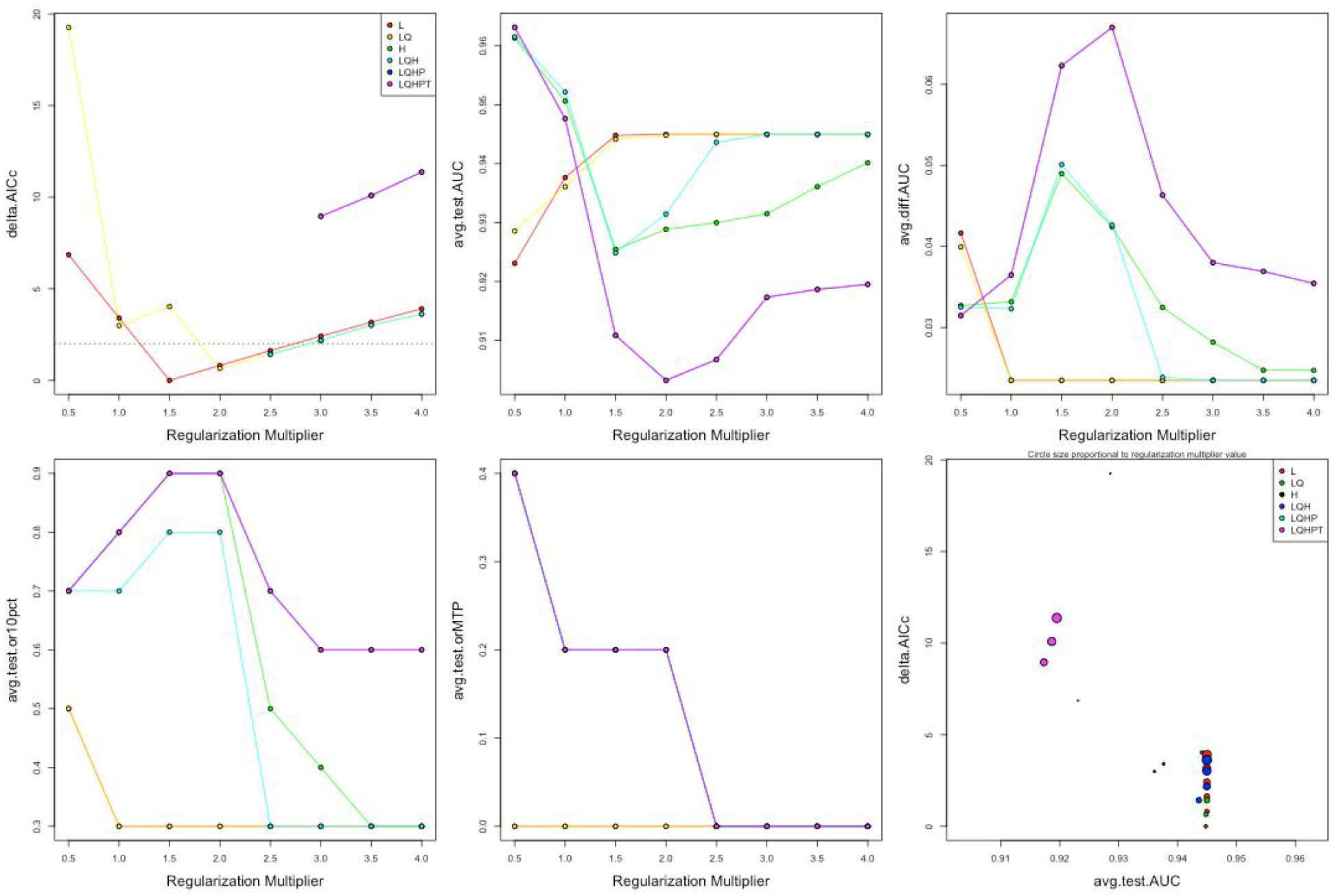
Hyperparameter tuning and model evaluation of *Z. clava-herculis* distribution during the T1 period.

**Sup. Fig. 6:**
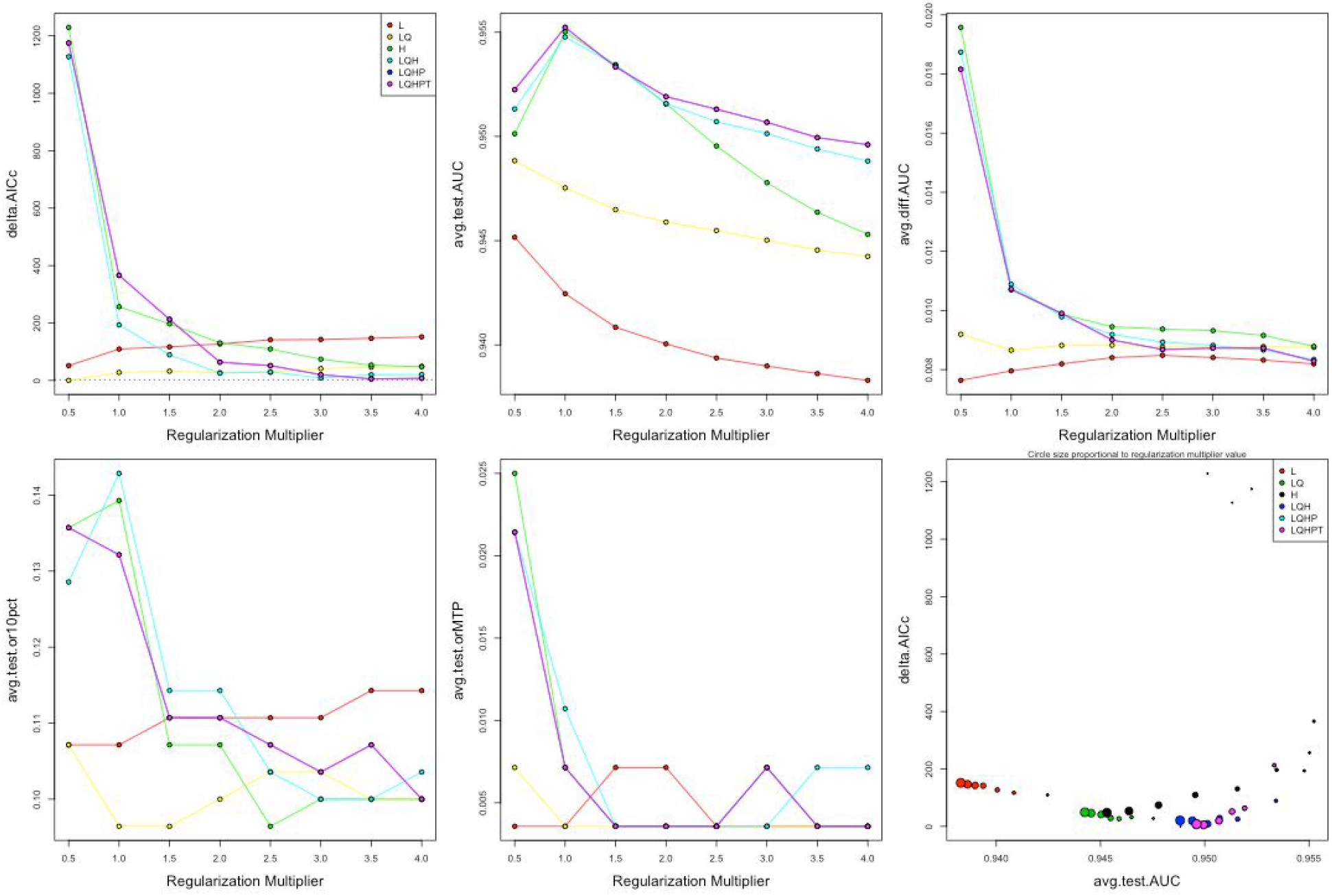
Hyperparameter tuning and model evaluation of *Z. clava-herculis* distribution during the T2 period.

**Sup. Fig. 7:**
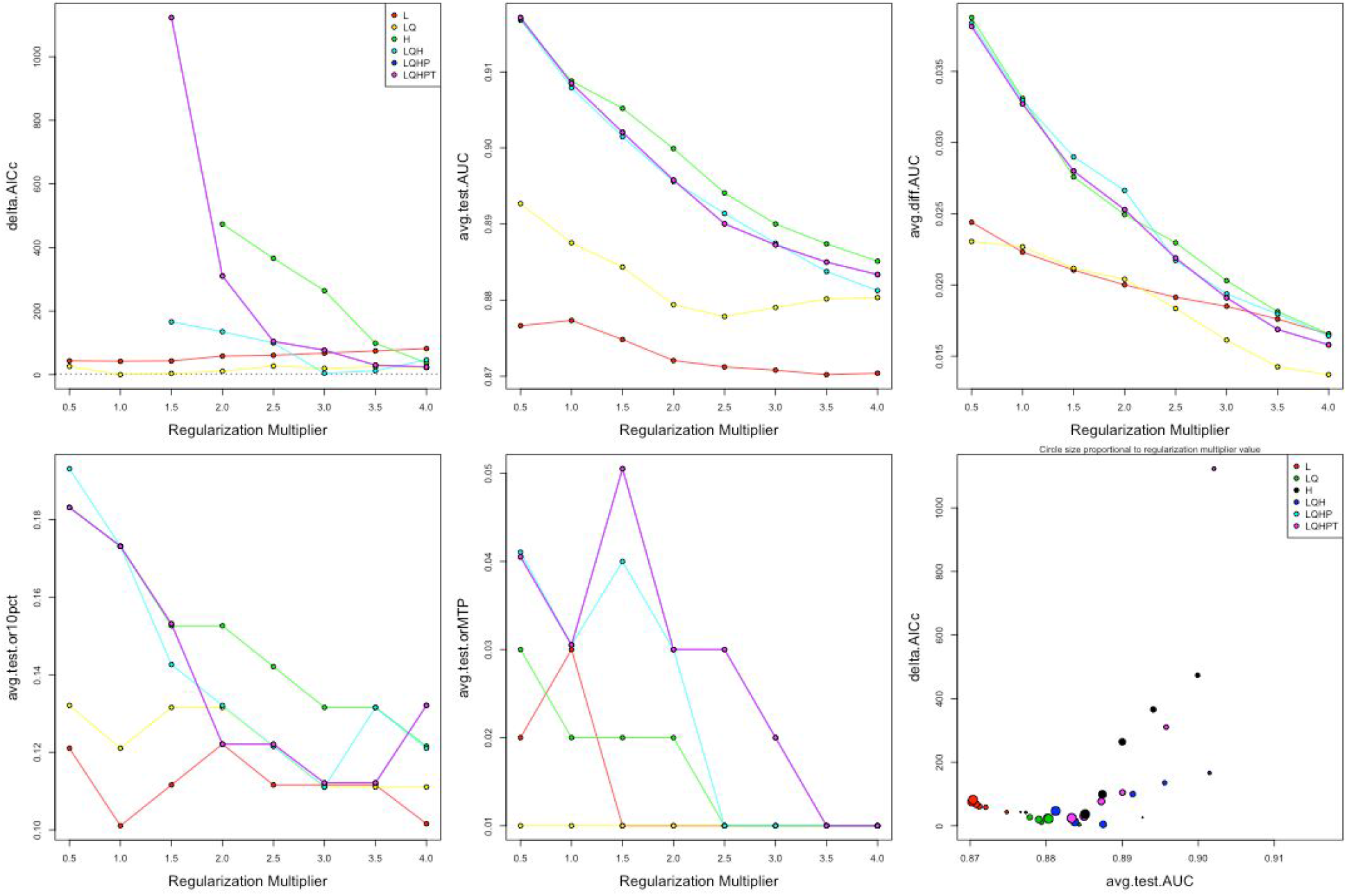
Hyperparameter tuning and model evaluation of *P. trifoliata* distribution during the T1 period.

**Sup. Fig. 8:**
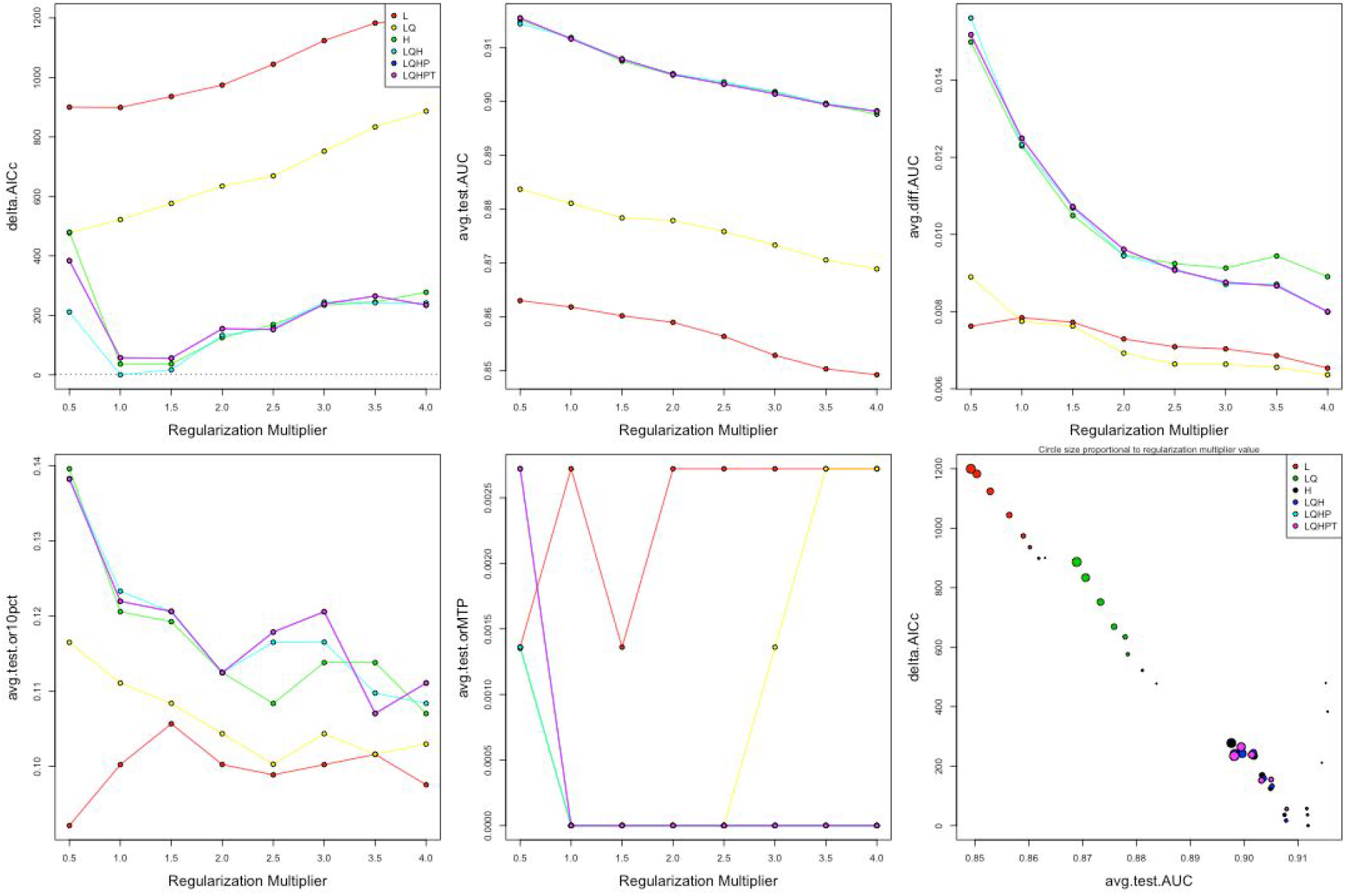
Hyperparameter tuning and model evaluation of *P. trifoliata* distribution during the T2 period.

**Supplementary Table 1.** Bioclim variable definitions

| **Bioclim Number** | **Definition** |
| --- | --- |
| BIO1 | Annual mean temperature |
| BIO2 | Mean diurnal range |
| BIO3 | Isothermality |
| BIO4 | Temperature seasonality |
| BIO5 | Max temperature of warmest month |
| BIO6 | Min temperature of warmest month |
| BIO7 | Temperature annual range |
| BIO8 | Mean temperature wettest quarter |
| BIO9 | Mean temperature driest quarter |
| BIO10 | Mean temperature of warmest quarter |
| BIO11 | Mean temperature of coldest quarter |
| BIO12 | Annual precipitation |
| BIO13 | Precipitation of wettest month |
| BIO14 | Precipitation of driest month |
| BIO15 | Precipitation seasonality |
| BIO16 | Precipitation of wettest quarter |
| BIO17 | Precipitation of driest quarter |
| BIO18 | Precipitation of warmest quarter |
| BIO19 | Precipitation of coldest quarter |

## References

Abatzoglou, J.T., Dobrowski, S.Z., Parks, S.A. & Hegewisch, K.C. (2018) TerraClimate, a high-resolution global dataset of monthly climate and climatic water balance from 1958-2015. Scientific data, 5, 170191.

Altieri, M.A., Nicholls, C.I., Henao, A. & Lana, M.A. (2015) Agroecology and the design of climate change-resilient farming systems. Agronomy for Sustainable Development, 35, 869–890.

Audusseau, H., Le Vaillant, M., Janz, N., Nylin, S., Karlsson, B. & Schmucki, R. (2017) Species range expansion constrains the ecological niches of resident butterflies. Journal of Biogeography, 44, 28–38.

Bader, M.Y., van Geloof, I. & Rietkerk, M. (2007) High solar radiation hinders tree regeneration above the alpine treeline in northern Ecuador. Plant Ecology, 191, 33–45.

Bale, J.S., Masters, G.J., Hodkinson, I.D., Awmack, C., Martijn Bezemer, T., Brown, V.K., Butterfield, J., Buse, A., Coulson, J.C., Farrar, J., Good, J.E.G., Harrington, R., Hartley, S., Hefin Jones, T., Lindroth, R.L., Malcolm C. Press, Symrnioudis, I., Watt, A.D. & Whittaker, J.B. (2002) Herbivory in global climate change research: direct effects of rising temperature on insect herbivores. Global Change Biology, 8, 1–16.

Battisti, A., Stastny, M., Netherer, S., Robinet, C., Schopf, A., Roques, A. & Larsson, S. (2005) Expansion of geographic range in the Pine Processionary Moth caused by increased winter temperatures. Ecological Applications, 15, 2084–2096.

Blois, J.L., Zarnetske, P.L., Fitzpatrick, M.C. & Finnegan, S. (2013) Climate change and the past, present, and future of biotic interactions. Science, 341, 499–504.

Bonney, R., Cooper, C.B., Dickinson, J., Kelling, S., Phillips, T., Rosenberg, K.V. & Shirk, J. (2009) Citizen Science: A Developing Tool for Expanding Science Knowledge and Scientific Literacy. BioScience, 59, 977–984.

Bonney, R., Shirk, J.L., Phillips, T.B., Wiggins, A., Ballard, H.L., Miller-Rushing, A.J. & Parrish, J.K. (2014) Citizen science. Next steps for citizen science. Science, 343, 1436–1437.

Breed, G.A., Stichter, S. & Crone, E.E. (2012) Climate-driven changes in northeastern US butterfly communities. Nature climate change, 3, 142.

Brown, C.J., O’Connor, M.I., Poloczanska, E.S., Schoeman, D.S., Buckley, L.B., Burrows, M.T., Duarte, C.M., Halpern, B.S., Pandolfi, J.M., Parmesan, C. & Richardson, A.J. (2016) Ecological and methodological drivers of species’ distribution and phenology responses to climate change. Global change biology, 22, 1548–1560.

Buckley, R., Morrison, C., Guy Castley, J., de Vasconcellos Pegas, F. & Mossaz, A. (2013) Conservation Science and Practice. Biological Conservation, 162, 133.

Bueno de Mesquita, C.P., King, A.J., Schmidt, S.K., Farrer, E.C. & Suding, K.N. (2016) Incorporating biotic factors in species distribution modeling: are interactions with soil microbes important? Ecography, 39, 970–980.

Cannon, R.J.C. (1998) The implications of predicted climate change for insect pests in the UK, with emphasis on non-indigenous species. Global Change Biology, 4, 785–796.

Castex, V., Beniston, M., Calanca, P., Fleury, D. & Moreau, J. (2018) Pest management under climate change: The importance of understanding tritrophic relations. The Science of the total environment, 616-617, 397–407.

Chalcoff, V.R., Aizen, M.A. & Ezcurra, C. (2012) Erosion of a pollination mutualism along an environmental gradient in a south Andean treelet, Embothrium coccineum (Proteaceae). Oikos, 121, 471–480.

Chamberlain, S., Ram, K. & Hart, T. (2016) spocc: Interface to species occurrence data sources. R package version 0. 5. 0. Website (https://CRAN.R-project.org/package$1/4$spocc).

Connell, J.H. (1961) The Influence of Interspecific Competition and Other Factors on the Distribution of the Barnacle Chthamalus Stellatus. Ecology, 42, 710–723.

Dempster, J.P. & Pollard, E. (1981) Fluctuations in resource availability and insect populations. Oecologia, 50, 412–416.

Descombes, P., Pradervand, J.-N., Golay, J., Guisan, A. & Pellissier, L. (2016) Simulated shifts in trophic niche breadth modulate range loss of alpine butterflies under climate change. Ecography, 39, 796–804.

Deutsch, C.A., Tewksbury, J.J., Huey, R.B., Sheldon, K.S., Ghalambor, C.K., Haak, D.C. & Martin, P.R. (2008) Impacts of climate warming on terrestrial ectotherms across latitude. Proceedings of the National Academy of Sciences of the United States of America, 105, 6668–6672.

Dickinson, J.L., Zuckerberg, B. & Bonter, D.N. (2010) Citizen Science as an Ecological Research Tool: Challenges and Benefits. Annual Review of Ecology, Evolution, and Systematics, 41, 149–172.

Dilts, T.E., Steele, M.O., Engler, J.D., Pelton, E.M., Jepsen, S.J., McKnight, S.J., Taylor, A.R., Fallon, C.E., Black, S.H., Cruz, E.E., Craver, D.R. & Forister, M.L. (2019) Host Plants and Climate Structure Habitat Associations of the Western Monarch Butterfly. Frontiers in Ecology and Evolution, 7.

Elith, J. & Leathwick, J.R. (2009) Species Distribution Models: Ecological Explanation and Prediction Across Space and Time. Annual Review of Ecology, Evolution, and Systematics, 40, 677–697.

Elith, J., Phillips, S.J., Hastie, T., Dudík, M., Chee, Y.E. & Yates, C.J. (2011) A statistical explanation of MaxEnt for ecologists. Diversity and Distributions, 17, 43–57.

Ettinger, A. & HilleRisLambers, J. (2017) Competition and facilitation may lead to asymmetric range shift dynamics with climate change. Global change biology, 23, 3921–3933.

Farquhar, Donald W. 1934. The Lepidoptera of New England. Harvard University, Ph.D. dissertation.

Fick, S.E. & Hijmans, R.J. (2017) WorldClim 2: new 1-km spatial resolution climate surfaces for global land areas. International Journal of Climatology, 37, 4302–4315.

Finkbeiner, S.D., Reed, R.D., Dirig, R. & Losey, J.E. (2011) The Role of Environmental Factors in the Northeastern Range Expansion ofPapilio cresphontesCramer (Papilionidae). Journal of the Lepidopterists’ Society, 65, 119–125.

Freeman, R.S., Brody, A.K. & Neefus, C.D. (2003) Flowering phenology and compensation for herbivory in Ipomopsis aggregata. Oecologia, 136, 394–401.

Gallagher, R.V., Hughes, L. & Leishman, M.R. (2013) Species loss and gain in communities under future climate change: consequences for functional diversity. Ecography, 36, 531–540.

Harrington, R., Fleming, R.A. & Woiwod, I.P. (2001) Climate change impacts on insect management and conservation in temperate regions: can they be predicted? Agricultural and Forest Entomology, 3, 233–240.

Hickling, R., Roy, D.B., Hill, J.K., Fox, R. & Thomas, C.D. (2006) The distributions of a wide range of taxonomic groups are expanding polewards. Global Change Biology, 12, 450–455.

Hijmans, R.J., Phillips, S., Leathwick, J., Elith, J. & Hijmans, M.R.J. (2017) Package “dismo.” Circles, 9, 1–68.

HilleRisLambers, J., Harsch, M.A., Ettinger, A.K., Ford, K.R. & Theobald, E.J. (2013) How will biotic interactions influence climate change-induced range shifts? Annals of the New York Academy of Sciences, 1297, 112–125.

Holt, R.D. (2003) On the evolutionary ecology of species’ ranges. Evolutionary ecology research, 5, 159–178.

Huey, R.B., Deutsch, C.A., Tewksbury, J.J., Vitt, L.J., Hertz, P.E., Álvarez Pérez, H.J. & Garland, T. (2009) Why tropical forest lizards are vulnerable to climate warming. Proceedings of the Royal Society B: Biological Sciences, 276, 1939–1948.

Kerr, J.T., Pindar, A., Galpern, P., Packer, L., Potts, S.G., Roberts, S.M., Rasmont, P., Schweiger, O., Colla, S.R., Richardson, L.L., Wagner, D.L., Gall, L.F., Sikes, D.S. & Pantoja, A. (2015) CLIMATE CHANGE. Climate change impacts on bumblebees converge across continents. Science, 349, 177–180.

Kéry, M., Gardner, B. & Monnerat, C. (2010) Predicting species distributions from checklist data using site-occupancy models. Journal of Biogeography, no-no.

Kingsolver, J.G., Woods, H.A., Buckley, L.B., Potter, K.A., MacLean, H.J. & Higgins, J.K. (2011) Complex life cycles and the responses of insects to climate change. Integrative and comparative biology, 51, 719–732.

Kosmala, M., Wiggins, A., Swanson, A. & Simmons, B. (2016) Assessing data quality in citizen science. Frontiers in Ecology and the Environment, 14, 551–560.

Lany, N.K., Zarnetske, P.L., Schliep, E.M., Schaeffer, R.N., Orians, C.M., Orwig, D.A. & Preisser, E.L. (2018) Asymmetric biotic interactions and abiotic niche differences revealed by a dynamic joint species distribution model. Ecology, 99, 1018–1023.

Lemoine, N.P. (2015) Climate change may alter breeding ground distributions of eastern migratory monarchs (Danaus plexippus) via range expansion of Asclepias host plants. PloS one, 10, e0118614.

Leroux, S.J., Larrivée, M., Boucher-Lalonde, V., Hurford, A., Zuloaga, J., Kerr, J.T. & Lutscher, F. (2013) Mechanistic models for the spatial spread of species under climate change. Ecological applications: a publication of the Ecological Society of America, 23, 815–828.

Liu, C., Berry, P.M., Dawson, T.P. & Pearson, R.G. (2005) Selecting thresholds of occurrence in the prediction of species distributions. Ecography, 28, 385–393.

Losey, J.E. & Vaughan, M. (2006) The Economic Value of Ecological Services Provided by Insects. BioScience, 56, 311.

Louthan, A.M., Doak, D.F. & Angert, A.L. (2015) Where and When do Species Interactions Set Range Limits? Trends in Ecology & Evolution, 30, 780–792.

Marlon, J.R., Pederson, N., Nolan, C., Goring, S., Shuman, B., Booth, R., Bartlein, P.J., Berke, M.A., Clifford, M., Cook, E., Dieffenbacher-Krall, A., Dietze, M.C., Hessl, A., Bradford Hubeny, J., Jackson, S.T., Marsicek, J., McLachlan, J., Mock, C.J., Moore, D.J.P., Nichols, J., Robertson, A., Schaefer, K., Trouet, V., Umbanhowar, C., Williams, J.W. & Yu, Z. (2016) Climatic history of the northeastern United States during the past 3000 years. Climate of the Past Discussions, 1–38.

Moeller, D.A., Geber, M.A., Eckhart, V.M. & Tiffin, P. (2012) Reduced pollinator service and elevated pollen limitation at the geographic range limit of an annual plant. Ecology, 93, 1036–1048.

Morales, N.S., Fernández, I.C. & Baca-González, V. (2017) MaxEnt’s parameter configuration and small samples: are we paying attention to recommendations? A systematic review. PeerJ, 5, e3093.

Muscarella, R., Galante, P.J., Soley-Guardia, M., Boria, R.A., Kass, J.M., Uriarte, M. & Anderson, R.P. (2014) ENMeval: An R package for conducting spatially independent evaluations and estimating optimal model complexity forMaxentecological niche models. Methods in Ecology and Evolution, 5, 1198–1205.

Palacio, F.X. & Girini, J.M. (2018) Biotic interactions in species distribution models enhance model performance and shed light on natural history of rare birds: a case study using the straight-billed reedhaunter Limnoctites rectirostris. Journal of Avian Biology, 49, e01743.

Parmesan, C., Ryrholm, N., Stefanescu, C., Hill, J.K., Thomas, C.D., Descimon, H., Huntley, B., Kaila, L., Kullberg, J., Tammaru, T., John Tennent, W., Thomas, J.A. & Warren, M. (1999) Poleward shifts in geographical ranges of butterfly species associated with regional warming. Nature, 399, 579–583.

Parmesan, C. & Yohe, G. (2003) A globally coherent fingerprint of climate change impacts across natural systems. Nature, 421, 37–42.

Pearson, D.L. & Knisley, C.B. (1985) Evidence for Food as a Limiting Resource in the Life Cycle of Tiger Beetles (Coleoptera: Cicindelidae). Oikos, 45, 161.

Phillips, S.J., Anderson, R.P., Dudík, M., Schapire, R.E. & Blair, M.E. (2017) Opening the black box: an open-source release of Maxent. Ecography, 40, 887–893.

Phillips, S.J., Anderson, R.P. & Schapire, R.E. (2006) Maximum entropy modeling of species geographic distributions. Ecological Modelling, 190, 231–259.

Phillips, S.J. & Dudík, M. (2008) Modeling of species distributions with Maxent: new extensions and a comprehensive evaluation. Ecography, 0, 080328142746259–???

Poloczanska, E.S., Brown, C.J., Sydeman, W.J., Kiessling, W., Schoeman, D.S., Moore, P.J., Brander, K., Bruno, J.F., Buckley, L.B., Burrows, M.T., Duarte, C.M., Halpern, B.S., Holding, J., Kappel, C.V., O’Connor, M.I., Pandolfi, J.M., Parmesan, C., Schwing, F., Thompson, S.A. & Richardson, A.J. (2013) Global imprint of climate change on marine life. Nature Climate Change, 3, 919–925.

Porter, J.H., Parry, M.L. & Carter, T.R. (1991) The potential effects of climatic change on agricultural insect pests. Agricultural and Forest Meteorology, 57, 221–240.

Pöyry, J., Luoto, M., Heikkinen, R.K., Kuussaari, M. & Saarinen, K. (2009) Species traits explain recent range shifts of Finnish butterflies. Global Change Biology, 15, 732–743.

Radosavljevic, A. & Anderson, R.P. (2014) Making better Maxentmodels of species distributions: complexity, overfitting and evaluation. Journal of Biogeography, 41, 629–643.

Rahel, F.J. & Olden, J.D. (2008) Assessing the effects of climate change on aquatic invasive species. Conservation biology: the journal of the Society for Conservation Biology, 22, 521–533.

Robillard, C.M., Coristine, L.E., Soares, R.N. & Kerr, J.T. (2015) Facilitating climate-change-induced range shifts across continental land-use barriers. Conservation biology: the journal of the Society for Conservation Biology, 29, 1586–1595.

Robinet, C. & Roques, A. (2010) Direct impacts of recent climate warming on insect populations. Integrative zoology, 5, 132–142.

Robinson, A., Inouye, D.W., Ogilvie, J.E. & Mooney, E.H. (2017) Multitrophic interactions mediate the effects of climate change on herbivore abundance. Oecologia, 185, 181–190.

Roth, T., Plattner, M. & Amrhein, V. (2014) Plants, birds and butterflies: short-term responses of species communities to climate warming vary by taxon and with altitude. PloS one, 9, e82490.

Sánchez-Bayo, F. & Wyckhuys, K.A.G. (2019) Worldwide decline of the entomofauna: A review of its drivers. Biological Conservation, 232, 8–27.

Scudder, S.H. (1889) The butterflies of the eastern United States and Canada with special reference to New England. Vol. II. Cambridge [Mass.] :S.H. Scudder. pp. 767–1774

Sexton, J.P., McIntyre, P.J., Angert, A.L. & Rice, K.J. (2009) Evolution and Ecology of Species Range Limits. Annual Review of Ecology, Evolution, and Systematics, 40, 415–436.

Speed, J.D.M., Austrheim, G., Hester, A.J. & Mysterud, A. (2010) Experimental evidence for herbivore limitation of the treeline. Ecology, 91, 3414–3420.

Stanton-Geddes, J., Tiffin, P. & Shaw, R.G. (2012) Role of climate and competitors in limiting fitness across range edges of an annual plant. Ecology, 93, 1604–1613.

Stueve, K.M., Isaacs, R.E., Tyrrell, L.E. & Densmore, R.V. (2011) Spatial variability of biotic and abiotic tree establishment constraints across a treeline ecotone in the Alaska Range. Ecology, 92, 496–506.

Svancara, L.K., Abatzoglou, J.T. & Waterbury, B. (2019) Modeling Current and Future Potential Distributions of Milkweeds and the Monarch Butterfly in Idaho. Frontiers in Ecology and Evolution, 7.

Swets, J. (1988) Measuring the accuracy of diagnostic systems. Science, 240, 1285–1293.

Urban, M.C., Bocedi, G., Hendry, A.P., Mihoub, J.-B., Pe’er, G., Singer, A., Bridle, J.R., Crozier, L.G., De Meester, L., Godsoe, W., Gonzalez, A., Hellmann, J.J., Holt, R.D., Huth, A., Johst, K., Krug, C.B., Leadley, P.W., Palmer, S.C.F., Pantel, J.H., Schmitz, A., Zollner, P.A. & Travis, J.M.J. (2016) Improving the forecast for biodiversity under climate change. Science, 353.

Urban, M.C., Phillips, B.L., Skelly, D.K. & Shine, R. (2007) The cane toad’s (Chaunus [Bufo] marinus) increasing ability to invade Australia is revealed by a dynamically updated range model. Proceedings. Biological sciences / The Royal Society, 274, 1413–1419.

Valavi, R., Elith, J., Lahoz-Monfort, J.J. & Guillera-Arroita, G. (2019a) blockCV: an R package for generating spatially or environmentally separated folds for k-fold cross-validation of species distribution models. Methods in Ecology and Evolution, 10, 225–232.

Valavi, R., Elith, J., Lahoz-Monfort, J.J. & Guillera-Arroita, G. (2019b) block CV: An r package for generating spatially or environmentally separated folds for k-fold cross-validation of species distribution models. Methods in Ecology and Evolution, 10, 225–232.

Warren, M.S., Hill, J.K., Thomas, J.A., Asher, J., Fox, R., Huntley, B., Roy, D.B., Telfer, M.G., Jeffcoate, S., Harding, P., Jeffcoate, G., Willis, S.G., Greatorex-Davies, J.N., Moss, D. & Thomas, C.D. (2001) Rapid responses of British butterflies to opposing forces of climate and habitat change. Nature, 414, 65–69.

Wisz, M.S., Hijmans, R.J., Li, J., Peterson, A.T., Graham, C.H., Guisan, A. & NCEAS Predicting Species Distributions Working Group† (2008) Effects of sample size on the performance of species distribution models. Diversity and Distributions, 14, 763–773.

Wisz, M.S., Pottier, J., Kissling, W.D., Pellissier, L., Lenoir, J., Damgaard, C.F., Dormann, C.F., Forchhammer, M.C., Grytnes, J.-A., Guisan, A., Heikkinen, R.K., Høye, T.T., Kühn, I., Luoto, M., Maiorano, L., Nilsson, M.-C., Normand, S., Öckinger, E., Schmidt, N.M., Termansen, M., Timmermann, A., Wardle, D.A., Aastrup, P. & Svenning, J.-C. (2013) The role of biotic interactions in shaping distributions and realised assemblages of species: implications for species distribution modelling. Biological reviews of the Cambridge Philosophical Society, 88, 15–30.

Ylioja, T., Roininen, H., Ayres, M.P., Rousi, M. & Price, P.W. (1999) Host-driven population dynamics in an herbivorous insect. Proceedings of the National Academy of Sciences of the United States of America, 96, 10735–10740.

Yu, J., Wong, W.-K. & Hutchinson, R.A. (2010) Modeling Experts and Novices in Citizen Science Data for Species Distribution Modeling. 2010 IEEE International Conference on Data Mining.

Zvereva, E.L., Hunter, M.D., Zverev, V. & Kozlov, M.V. (2016) Factors affecting population dynamics of leaf beetles in a subarctic region: The interplay between climate warming and pollution decline. Science of The Total Environment, 566–567, 1277–1288.

Battisti, A., Stastny, M., Netherer, S., Robinet, C., Schopf, A., Roques, A. & Larsson, S. (2005) Expnasion of geographic range in the pine processionary moth caused by increased winter temperatures. Ecological Applications, 15, 2084–2096.

Thompson, J. (1988) Variation in preference and specificity in monophagous and oligiophagous swallowtail butterflies. Evolution, 42, 118–218.

